# Nutrient Sensing Receptor GPRC6A Regulates mTORC1 Signaling and Tau Biology

**DOI:** 10.1101/2024.03.24.586459

**Authors:** Chao Ma, Kelsey Campbell, Andrii Kovalenko, Leslie A. Sandusky-Beltran, Huimin Liang, Jerry B. Hunt, John Calahatian, Mani Kallupurackal, Shalini Pandey, Muskan Vasisht, Mallory Watler, Zainuddin Quadri, Camilla Michalski, Margaret Fahnestock, Athanasios Papangelis, Daniel Sejer Pedersen, Trond Ulven, Kevin Nash, Maj-Linda B. Selenica, Dave Morgan, Paula C. Bickford, Daniel C. Lee

**Author notes:** **Corresponding Author:** Daniel C. Lee.

## Abstract

Tauopathies, including Alzheimer’s disease (AD), comprise microtubule-associated protein tau aggregates that cause neuronal cell death and clinical cognitive decline. Reducing overall tau abundance remains a central strategy for therapeutics; however, no disease-modifying treatment exists to date. One principal pathway for balancing cellular proteostasis includes the mechanistic target of rapamycin complex 1 (mTORC1) signaling. Recently, arginine emerged as one of the primary amino acids to activate mTORC1 through several intracellular arginine sensors and an extracellular arginine receptor, namely the G protein-coupled receptor (GPCR) family C, group 6, member A (GPRC6A). Human AD brains were previously reported with elevated mTORC1 signaling; however, it is unclear whether arginine sensing and signaling to mTORC1 plays a role in tauopathies. Herein, we examined arginine sensing associated with mTORC1 signaling in the human AD and animal models of tauopathy. We found that human AD brains maintained elevated levels of arginine sensors with potential uncoupling of arginine sensing pathways. Furthermore, we observed increased GPRC6A and arginine in the brain, accompanied by increased mTORC1 signaling and decreased autophagy in a mouse model of tauopathy (Tau PS19). We also discovered that both supplementing arginine and overexpressing GPRC6A in cell culture models could independently activate mTORC1 and promote tau accumulation. In addition, we found that suppressing GPRC6A signaling by either genetic reduction or pharmacological antagonism reduced tau accumulation, phosphorylation, and oligomerization. Overall, these findings uncover the crucial role of arginine sensing pathways in deregulating mTORC1 signaling in tauopathies and identify GPRC6A as a promising target for future therapeutics in tauopathies and other proteinopathies.

**Significance Statement:** Tauopathies, including Alzheimer’s disease (AD), accumulate pathogenic tau protein inclusions that potentially contribute to the hyperactive mechanistic target of rapamycin complex 1 (mTORC1) signaling and eventually cause neuronal cell death. Here, we presented novel findings that AD and animal models of tauopathy maintained increased expression of arginine sensors and uncoupling of arginine sensing associated with mTORC1 signaling. We investigated the role of a putative extracellular arginine and basic L-amino acid sensing G protein-coupled receptor (GPCR) family C, group 6, member A (GPRC6A) in activating mTORC1 and accelerating pathogenic tau phenotypes in several cell models. Additionally, we showed that genetic repression or antagonism of GPRC6A signaling provides a novel therapeutic target for tauopathies and other proteinopathies.

## Introduction

Alzheimer’s disease (AD) is mainly diagnosed with pathogenic proteins of intracellular neurofibrillary tangles (NFTs) and extracellular amyloid-β (Aβ) plaques (Masters et al., 2015). Tauopathies consist of a broader group of neurodegenerative diseases characterized by pathogenic protein inclusions formed by abnormal accumulation of microtubule-associated protein tau (MAPT) (Morris et al., 2011; Spillantini and Goedert, 2013). Recent AD positron emission tomography (PET) data shows that tau-PET predicts subsequent brain atrophy but not Aβ-PET, indicating that tau pathology is a primary driver of neurodegeneration in AD (La Joie et al., 2020). Among many anti-tau approaches, one common principle argues that reducing overall tau levels or preventing tau accumulation is critical to decreasing or even reversing tauopathies (DeVos et al., 2017; Lasagna-Reeves et al., 2016; Maeda and Mucke, 2016).

Considering that large tau inclusions become demanding clients for the ubiquitin-proteasome system, methods to improve the autophagy-lysosome system hold promise (Spillantini and Goedert, 2013). Increasing reports in AD patients indicate altered arginine and polyamine metabolism in AD, which possibly aggravates tau pathology by impairing proteostasis through signaling between the mechanistic target of rapamycin complex 1 (mTORC1) and autophagy (Graham et al., 2015; Liu et al., 2014; Morrison and Kish, 1995; Vemula et al., 2019). We have previously uncovered a unique association between arginine metabolism and mTORC1-autophagy signaling in AD mouse models of tau or Aβ independently (Hunt et al., 2015; Ma et al., 2021a; Ma et al., 2021b). Others have shown that stimuli in different animal models of AD can promote dysregulation of arginine metabolism, causing uncoupling and impaired clearance (Kan et al., 2015). Emerging evidence indicates that arginine is a necessary amino acid through its interaction with specific amino acid sensors to activate mTORC1. (Chantranupong et al., 2016; Rebsamen et al., 2015; Wolfson and Sabatini, 2017).

Although mTORC1 signaling is increased in AD brains, it is unknown if arginine sensing associated with mTORC1 plays a role in tau deposition (Mueed et al., 2018; Oddo, 2012; Pei and Hugon, 2008; Wang et al., 2014). Furthermore, recent fundamental studies revealed that G protein-coupled receptor (GPCR) family C, group 6, member A (GPRC6A), an extracellular arginine and basic L-amino acid receptor receptor, signals to mTORC1 signaling (Jacobsen et al., 2013; Wellendorph et al., 2005; Ye et al., 2017; Ye et al., 2019).

In the present study, we determined to understand if arginine sensing associated mTORC1 signaling was dysregulated in AD and a mouse tauopathy model and if manipulation of GRPC6A signaling impacts tau phenotypes. By profiling gene transcripts involved in arginine sensing mTORC1 signaling and assessing its key protein expression levels in AD and Tau PS19 mice, we found aberrant mTORC1 activation and uncoupling of arginine sensing pathway during tauopathies. Both *in vitro* and *in vivo* manipulation of the putative arginine sensor GPRC6A further validates it as a novel broad therapeutic target in treating diseases of mTORopathy and proteinopathy.

## Materials and Methods

### Experimental design and statistical analyses

Sample sizes of animal models and human patients were not predetermined since the effect sizes were unknown. At least five animals or eight human patients per group were used for the in vivo study, except for 1-2 animals to collect microdialysis data from 3-6 time points. All in vitro cell culture experiments were independently repeated 3-4 times with different biological replicates. No samples were excluded from the analysis. Mixed littermates from different inbred mice were used for assigning experimental groups based on genotype, age, and gender.

Detailed statistical analyses were summarized in Table 1 (Summary of Statistical Analysis). Statistical analysis and graphs were generated using SPSS (version 25.0, IBM Corp., Armonk, NY, USA) and GraphPad Prism (version 9.2.0, GraphPad Software, San Diego, CA, USA). The data met the standard distribution assumption with similar estimated variance among groups for the intended statistical tests. Comparisons between two groups or at least three groups were analyzed using a two-tailed unpaired Student’s t-test or one-way ANOVA with indicated post hoc multiple-comparison test, respectively. In the figures, all the data are presented as mean ± SEM. The mean difference was considered statistically significant at or below the p-value of 0.05.

**Table 1:**
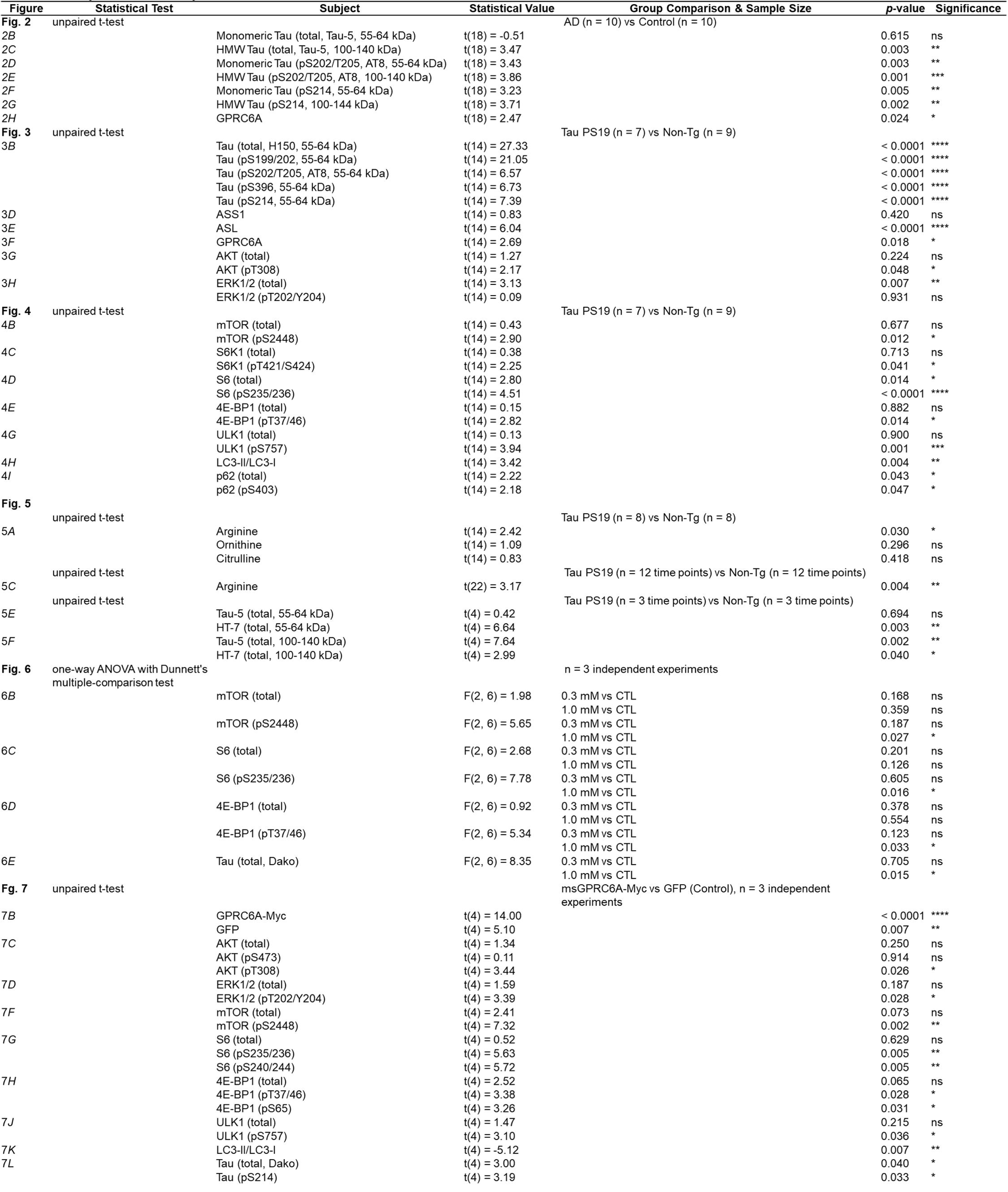
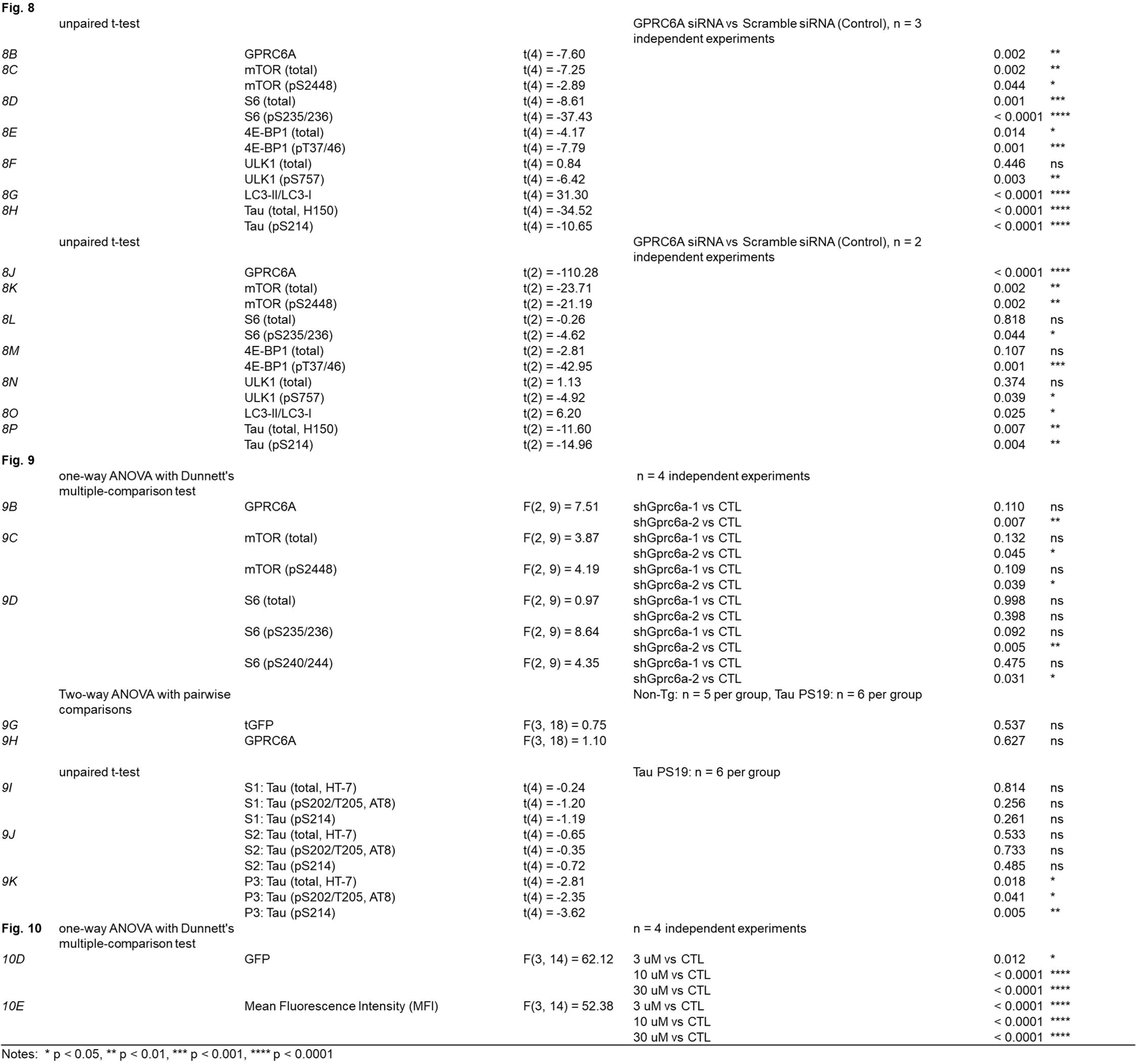
Summary of Statistical Analysis.

### Human Brain mRNA Extraction

As previously described, the total mRNA was extracted (Sandusky-Beltran et al., 2021). The Institute of Brain Aging and Dementia Tissue Repository provided human postmortem brain hippocampal tissues at the University of California, Irvine, CA, USA. A total of 16 samples were classified into two groups: Alzheimer’s disease patients (n = 8 including four males and four females, average age = 78, average PMI = 3.65 hours) and control patients (n = 8 including two males and six females, average age = 73, average PMI = 6.40 hours) based on matched age and gender. Alzheimer’s disease patients were confirmed by clinical and pathological diagnosis. All samples were collected at autopsy and snap frozen at -80 °C until use.

### Real-Time Quantitative Reverse Transcription PCR (qRT-PCR)

The extracted mRNA was reverse transcribed into cDNA and then used for the subsequent qRT-PCR assay. A Customed PCR array RT^2^ Profiler (Qiagen, 330171) was designed to be preloaded with primers to measure a list of arginine sensing related mTORC1 and autophagy signaling genes and housekeeping genes. According to the manufacturer’s instructions, the qRT-PCR was performed with RT2 SYBR^®^ Green qPCR Mastermix (Qiagen, 330529) using the BioRad Opticon 2. The table of threshold cycle (CT) values assigned with AD and control groups was uploaded to the Qiagen Online Data Analysis Center for analysis and graphing. Three housekeeping genes (ACTB, GUSB, GAPDH) were the normalization control. The delta-delta CT approach was used for creating fold change values. The delta CT was calculated by the gene of interest and the average of selected housekeeping genes, then delta-delta CT was processed (delta CT (experiment)-delta CT (control)). The *p*-values were calculated using an unpaired t-test of each gene’s replicate 2∧ (-Delta CT) values.

### Human Brain Protein Extraction

Human postmortem brain cortex tissues were obtained from the NIH NeuroBioBank and extracted for total protein as previously described (Sandusky-Beltran et al., 2021). Brain tissues were dissected and frozen at -80 °C until use. Samples include neuropathologically confirmed Alzheimer’s disease (AD) cases (n = 10 including six males and four females, average age = 83, average PMI = 10.67 hours) and normal aging matched control cases (n = 10 including five males and five females, average age = 79, average PMI = 15.11 hours). Frozen brain cortex tissues were cut, weighed (10-20 mg), and homogenized in sterile 1.5 ml tubes. Samples were lysed at 20% w/v in modified RIPA buffer (50 mM Tris-HCl, 150 mM NaCl, 0.5% Triton X-100, 1.0% NP-40, 1.0% SDS, 1% Sodium Deoxycholate), containing 1% v/v of protease inhibitor cocktail (Sigma-Aldrich, #P8340-5ML), phosphatase inhibitor cocktail 2 (Sigma-Aldrich, #P5726-5ML), phosphatase inhibitor cocktail 3 (Sigma-Aldrich, #P0044-5ML), and PMSF (phenylmethane sulfonyl fluoride, Sigma-Aldrich, #93482-50ML-F). Samples were homogenized with a motorized pestle, followed by sonication to obtain the whole protein lysate. Protein concentration was detected by the Pierce™ BCA Protein Assay Kit (Thermo Scientific™, #23225). Whole protein lysate was thus extracted and used for the subsequent western blots.

### Animal Models

Mouse colony maintenance and experimental procedures were performed according to the guidelines and protocols evaluated and approved by the Institutional Animal Care and Use Committee (IACUC) at the Byrd Alzheimer’s Center and Research Institute at the University of South Florida. Mouse experiments were strategically designed on the genotype, gender, and littermate by using non-transgenic (Non-Tg; The Jackson Laboratory, C57BL/6J, #000664) and Tau PS19 transgenic mice (Tg (Prnp-MAPT*P301S) PS19Vle, The Jackson Laboratory, #008169). The Tau PS19 mice gain function on the human gene of the microtubule-associated protein tau (MAPT) with P301S mutation. The human MAPT transgene maintained at least 5-fold higher expression than the mouse endogenous MAPT gene under the promoter of mouse prion protein PRNP (Yoshiyama et al., 2007). The Tau PS19 mice were maintained as the C57BL/6 hemizygous genetic background, while the non-transgenic littermates were used as the control.

### Mouse Brain Protein Extraction

Mouse brain tissue was harvested for protein extraction as previously described (Sandusky-Beltran et al., 2021). Frozen tissue was weighed and emersed in 20% w/v in modified RIPA buffer (50mM Tris, 150mM NaCl, 0.5% Triton x100, 1mM EGTA, 3% SDS, 1% Sodium Deoxycholate) with four protease or phosphatase inhibitors at 1% (Sigma-Aldrich, #P8340-5ML, #P5726-5ML, #P0044-5ML, #93482-50ML-F). Whole-cell lysates were obtained by homogenizing tissues with a motorized pestle. Two-thirds of the whole cell lysate was centrifuged at 40,000 x g for 30 minutes at 4 °C. The supernatant fraction was the detergent-soluble lysate. Protein concentration was determined by the standard Pierce™ BCA Protein Assay Kit (Thermo Scientific™, #23225).

Protein extraction involving the sarkosyl solution was processed according to a previously published method (Berger et al., 2007). Briefly, tissues were homogenized in homogenization buffer (50 mM Tris-HCl, 274 mM NaCl, 5 mM KCl) with the inhibitors above and centrifuged at a low speed of 13,000 xg for 15 minutes at 4 °C to obtain the total supernatant extracts and a pellet (P1). The protein concentration of total extracts was measured by the BCA Protein Assay Kit (Thermo Scientific™, #23225). The same mass of total extracts was taken for a high spin at 154,000 g for 20 minutes in 4 °C using an ultracentrifuge (Beckman, TLX-120K) to obtain the supernatant soluble fraction (S1) and a second pellet (P2). The P2 pellet was resuspended in solubilization buffer (10 mM Tris-HCl, 0.8 M NaCl, 10% Sucrose, 1 mM EGTA) with the above inhibitors and spun down at high speed of 154,000 xg for 20 minutes at 4 °C to obtain the supernatant fraction which was further dissolved in 1% sarkosyl solution (Teknova, #S3376,) at 37 °C for 1 hour. Afterward, it’s spun down at high speed of 154,000 xg for 35 minutes at 4 °C to obtain the final supernatant fraction (sarkosyl soluble, S2) and the final pellet. The final pellet was dissolved in 1x TE buffer (Alfa Aesar, #J60738-AK), which became the sarkosyl insoluble fraction (P3).

### Western Blot

We performed western blot analysis as previously described using an Azure c600 imager (Azure Biosystems) (Sandusky-Beltran et al., 2021). Protein was loaded onto a 4∼20% Tris-Glycine gradient gels, transferred onto PVDF membrane blots, incubated first with an unconjugated primary antibody overnight at 4 °C, then with a secondary antibody conjugated with horseradish peroxidase (HRP), and finally detected by chemiluminescence after incubating with HRP substrate (Thermo Scientific™, #P32106). For mouse brain proteins, 2.5-5 µg was loaded to measure tau epitopes and 10-20 µg was loaded for other probing targets. For cell culture protein lysate, 20-30 µg was loaded. All groups were subjected to the same gel (resolving, transferring, and exposing) for a given probing target. Representative bands from all groups were cropped by Adobe Photoshop CS5 and used for densitometry analysis by AlphaEaseFCTM Software (Version 3.1.2, Alpha Innotech Corporation). Primary antibodies used in this study include Tau-5 (NeoMarkers, #MS-247-PABX), HT-7 (Thermo Scientific™, # MN1000), Tau pS202/T205 (AT8, Thermo Scientific™, #MN1020), Tau pS214 (abcam, #ab170892), Tau-H150 (Santa Cruz Biotechnology, #sc-5587), Tau PS199/202 (AnaSpec, #54963-025), Tau pS396 (AnaSpec, #54977-025), ASS1 (Protein Tech, #16210-1-AP), ASL (Protein Tech, #16645-1-AP), GPRC6A (human, Gene Tex, #GTX108214), GPRC6A (mouse, LifeSpan BioSciences, #LS-B14489), AKT (Cell Signaling Tech, #4691X), AKT pT308 (Cell Signaling Tech, #2965S), AKT pS473 (Cell Signaling Tech, #4060X), ERK1/2 (Cell Signaling Tech, #9102S), ERK1/2 pT202/Y204 (Cell Signaling Tech, #9101), mTOR (Cell Signaling Tech, #2983S), mTOR pS2448 (Cell Signaling Tech, #5536S), S6K1 (Cell Signaling Tech, #2708), S6K1 pT421/S424 (Cell Signaling Tech, #9204S), S6 (Cell Signaling Tech, #2217S), S6 pS235/236 (Cell Signaling Tech, #2211S), S6 pS240/244 (Cell Signaling Tech, #2215S), 4E-BP1 (Cell Signaling Tech, #9644), 4E-BP1 pT37/46 (Cell Signaling Tech, #2855), 4E-BP1 pS65 (Cell Signaling Tech, #9451S), ULK1 (Cell Signaling Tech, #8054), ULK1 pS757 (Cell Signaling Tech, #14202), p62 (Protein Tech, #18420-1-AP), p62 pS403 (Cell Signaling Tech, #39786S), LC3 (Cell Signaling Tech, #L7543), Tau Dako (Agilent Technologies, #A002401-2), C-Myc (Cell Signaling Tech, #2272), GFP (abcam, #ab13970), tGFP (turbo GFP, Origene, #TA150075), GAPDH (Meridian Life Science, #H86504M), and β-actin (Sigma Aldrich, #A5441). Secondary antibodies include Goat Anti-Rabbit IgG-HRP (Southern Biotech, #4010-05), Goat Anti-Mouse IgGI-HRP (Southern Biotech, #1070-05), and Goat Anti-Chicken IgG-HRP (Southern Biotech, # 6100-05). AzureRed Fluorescent Total Protein Stain (Azure Biosystems, #AC2124) was used for whole protein staining.

### In Vivo Microdialysis

In vivo microdialysis was performed to assess the brain interstitial fluid (ISF) tau and arginine levels from awake and freely moving 8.5-9.5-month-old non-transgenic and Tau PS19 mice as previously described (Yamada et al., 2014). Briefly, a guide cannula (Eicom/ Amuza) and dummy probe were stereotaxically implanted into the left hippocampus (LAT: -2.5, AP: -3.1, DV: -1.2) under isoflurane anesthesia and cemented. Immediately following recovery from surgery, dummy probes were replaced with 2-mm 1,000-kD cut-off AtmosLM microdialysis probes (Eicom/ Amuza) through the guide cannula. To account for the vent in the microdialysis probe, microdialysis perfusion was performed at a rate of 1.2 µl/min, and ISF was collected at a rate of 1.0 µl/min. Mice were given 24-hours to habituate to microdialysis perfusion and placement into RaTurn housing apparatuses (Bioanalytical Systems). As a perfusion buffer, 30% BSA (Sigma-Aldrich, #A9576-50ML) was diluted to 4% with artificial cerebrospinal fluid (aCSF; namely 1.3 mM CaCl_2_, 3 mM KCl, 0.4 mM KH^2^PO4, 1.2 mM MgSO_4_, 25 mM NaHCO_3_, 122 mM NaCl, and pH 7.35) on the day of use and filtered through a 0.1 µm membrane. Following the habituation period, baseline samples were collected before a 1-hour induction of neuronal activity by perfusion with 100 mM high K+ (97 mM KCl in aCSF was substituted for an equal amount of NaCl) by reverse dialysis. During sampling, ISF was continually collected in 1-hour increments and stored using a refrigerated fraction collector (SciPro). ISF collected from microdialysis was measured for tau by western blot and arginine by LC-MS/MS at the Sanford Burnham Prebys Institute (Orlando, FL, USA).

### Amino Acid Quantification

Total brain homogenates from 8.5∼9.5-month-old non-transgenic and Tau PS19 mice were harvested for liquid chromatography-mass spectrometry/mass spectrometry (LC-MS/MS) based on a standard curve of specific amino acid analytes. Arginine, Ornithine, and Citrulline quantification in brain homogenates was performed by Sanford Burnham Prebys (Orlando, FL, USA).

### Cell Cultures

All cell lines were maintained in T-flasks in a cell culture incubator (Forma 3110, Thermo Scientific) with 5% CO2 at 37 °C. Human iHEK-Tau cells were tetracycline-inducible human HEK-293 cells transiently expressing human wild type 4R0N tau. Human iHEK-Tau cells were cultured in a complete medium using DMEM (Gibco, #11965), supplemented with 10% heat-inactivated fetal bovine serum (HI-FBS, Sigma-Aldrich, #12306C), 1% GlutaMAX (200 mM, Gibco, #35050061), 1% penicillin (10,000 IU/ml)-streptomycin (10,000 µg/ml, Corning, #30-002-CI), 0.06% blasticidin (5 mg/ml, Invitrogen, #R210-01), and 0.125% zeocin (100 mg/ml, Invitrogen, #R25001). Human HeLa C3-Tau cells that were stably overexpressing human wild type 4R0N tau, were grown in a complete medium using Opti-MEM (Gibco, #31985), supplemented with 10% HI-FBS (Sigma-Aldrich, #12306C), 2% GlutaMAX (200 mM, Gibco, #35050061), 1% penicillin (10,000 IU/ml)-streptomycin (10,000 µg/ml, Corning, #30-002-CI), and 1% geneticin (50 mg/ml, Corning, #30234CI). Mouse naïve N2a cells (ATCC® CCL131TM) were cultured in DMEM (Gibco, #11965) supplemented with 10% HI-FBS (Sigma-Aldrich, #12306C), 1% GlutaMAX (200 mM, Gibco, #35050061), 1% penicillin (10,000 IU/ml)-streptomycin (10,000 µg/ml, Corning, #30-002-CI), and 1% sodium pyruvate (100 mM, Gibco, #11360070). E18 wild-type mouse primary cortical neurons were extracted as previously described and cultured in neurobasal medium (Invitrogen, #21103049) supplemented with 2% B-27 supplement (Invitrogen, #17504044) and 2% GlutaMAX (200 mM, Gibco, #35050061) (Woo et al., 2017). L-arginine (Sigma-Aldrich, #A5006-100G) was added in a concentration-dependent manner to the fourth day in vitro culture for 24 hours. For all cell culture experiments, whole-cell lysates were extracted using Mem-PER™ Plus Membrane Protein Extraction Kit (Thermo Scientific™, #89842) according to the manufacturer’s protocols. Protein concentration was assessed by the Pierce™ BCA Protein Assay Kit (Thermo Scientific™, #23225) for western blots.

### Cloning and Transfection of GPRC6A Plasmids

Mouse Gprc6a fused with Myc tag was cloned into the AAV backbone vector to generate a recombinant plasmid intended for future rAAV production. Briefly, the donor plasmid msGPRC6A-Myc (6812 bp) was amplified by PCR using primers of Ms.GPRC6A-Myc FWD (5’-GAATTTACCGGTGCCACCATGGTCCTTCTGTTGATC-3’) and Ms.GPRC6A-Myc REV (5’-GGGCCCGCAATTGTCATATACTTGAACTTCTTTTC-3’) to generate a sequence size of 2867 bp. The amplified PCR product was then digested by restriction enzymes of AgeI (New England Biolabs, #R0552S) and MfeI (New England Biolabs, #R0589S) to produce the insert. Meanwhile, the same enzymes cut the AAV virus backbone plasmid pTR-Up-MCSW (4697 bp) to create the vector. The insert fragment was thus ligated with the vector to produce the final product (7500 bp). The outcome of pTR-msGPRC6A-Myc recombinant plasmid expresses full-length mouse GPRC6A fused with the Myc tag under the promoter of Ubq C. A control plasmid pTR-Up-GFP-MCSW expresses GFP in the same vector backbone. The GFP plasmid and GPRC6A-Myc plasmid were transfected into the HEK293T that stably expresses a tetracycline-inducible wild-type human tau (4R0N) (iHEK-Tau) using Lipofectamine™ 2000 Transfection Reagent (Invitrogen™, #11668019). Doxycycline was added to the cells to induce tau production on the first day of transfection. Post 72-hour transfection, cells were harvested and extracted with whole-cell lysate for western blot analysis.

### Transfection of GPRC6A siRNAs and shRNAs

Predesigned siGENOME siRNAs against human GPRC6A (Horizon Discovery, #M-005613-01-0005) were purchased in SMARTpool format. The siGENOME Non-Targeting Control siRNAs (Horizon Discovery, #D-001210-03-05) were used as the scramble control siRNA to target no known genes in human cells. The Lipofectamine™ RNAiMAX Transfection Reagent (Invitrogen™, #13778150) was used for transfecting siRNAs into human HeLa C3-Tau cells that stably express human wild-type 4R0N tau and iHEK-Tau cells. Cells were harvested for whole protein extraction after 72 hours.

The pGFP-A-shAAV shRNA cloning plasmids against mouse Gprc6a (Origene, HC141118) were designed for transfection in mouse N2a cells and production for rAAVs. The pGFP-A-shAAV shRNA plasmid (Origene, #TR30034) served as the scramble control to produce turbo GFP (tGFP) as the reporter protein. The shGprc6a-1 shRNA (ATTGGTCAACCGCTACCAAGATTATCACC) and shGprc6a-2 shRNA (AAGACACCTGTCGTGAAATCATCTGGAGG) were cloned into the pGFP-A-shAAV shRNA cloning plasmid to produce shRNA and tGFP. All three plasmids were subsequently transfected into mouse N2a cells for western blot analysis.

### In Vitro Production of rAAVs

The pGFP-A-shAAV transfer plasmid (ITR-U6-scramble shRNA-CMV-tGFP-ITR) and pGFP-A-shGprc6a-2-shAAV transfer plasmid (ITR-U6-Gprc6a shRNA-2-CMV-tGFP-ITR) were individually co-transfected with packaging plasmid pAAV9 (Rep-Cap) and adenoviral helper plasmid pXX6 (pHelper) into HEK-293T cells for in vitro production of the complete viruses of rAAV-shScramble and rAAV-shGprc6a-2 according to the addgene protocol (AAV Production in HEK293T Cells). The rAAVs were then purified according to the addgene protocol (AAV Purification by Iodixanol Gradient Ultracentrifugation). Finally, physical titers of rAAV samples (viral genomes (vg)/ml) were measured by qPCR using the primers of hCMV-FWD (5’-GATGCGGTTTTGGCAGTACAT-3’) and hCMV-REV (5’-TTTTGGAAAGTCCCGTTGATTT-3’) according to the addgene protocol (AAV Titration by qPCR Using SYBR Green Technology) (Aurnhammer et al., 2012).

### In Vivo Injection of rAAVs

The standard stereotaxic surgical procedures for injecting viruses were performed as previously described (Sandusky-Beltran et al., 2019; Sandusky-Beltran et al., 2021). Briefly, the rAAV-shScramble and rAAV-shGprc6a-2 were bilaterally injected into 6-month-old Tau PS19 and Non-Tg mice with the same amount of virus (2 µl volume of 1.13E+13 vg/ml concentration) using a convection-enhanced delivery approach at the rate of 1.5 µl/ml. Viruses were injected into the anterior cortex (ACX, AP = -2.2 mm, ML = ±2.2 mm, DV = -2.5 mm), hippocampus (HPC, AP = 2.5 mm, ML = ±2.5 mm, DV = -2.7 mm), and entorhinal cortex (ECX, AP = 3.2 mm, ML = ±2.9 mm, DV = -5.0 mm). The rAAV viral constructs were incubated for up to 4 months, followed by brain tissue collection.

### Split GFP-Tau Oligomerization Assay

As previously described, the N2a split superfolder GFP-Tau (N2a-ssGT) cell line was used to measure tau oligomerization (Sandusky-Beltran et al., 2019; Sandusky-Beltran et al., 2021). The N2a-ssGT cells were cultured in the DMEM medium (Gibco, #11965) supplemented with 10% HI-FBS (Sigma-Aldrich, #12306C), 1% GlutaMAX (200 mM, Gibco, #35050061), 1% penicillin (10,000 IU/ml)-streptomycin (10,000 µg/ml, Corning, #30-002-CI), 1% sodium pyruvate (100 mM, Gibco, #11360070), zeocin (125 μg/ml, Life Technologies, #R25001), and geneticin (400 μg/ml, Corning, #30234CI) (Sandusky-Beltran et al., 2019; Sandusky-Beltran et al., 2021). Briefly, two plasmid constructs pmGFP10C-Tau (Addgene, #71433) and pmGFP11C-Tau (Addgene, #71434), were transfected into naïve N2a cells using Lipofectamine™ 2000 Transfection Reagent (Invitrogen™, #11668019). Each split GFP plasmid was fused with a human wild-type 4R0N monomer tau. The lead GPRC6A allosteric antagonist compound Cpd47661 (MW 391.47) was synthesized according to the published method and analytical data was in agreement with that reported for Cpd 3 (Gloriam et al., 2011) . The Cpd47661 was applied to the N2a-ssGT cells at 3 µM, 10 µM, and 30 µM for 72 hours. The exact amount of DMSO solvent vehicle served as the control. N2a-ssGT cells were imaged for GFP fluorescence using CytationTM 3 Cell Imaging Multi-Mode Reader (BioTekTM) and flow cytometry analysis by Accuri^®^ C6 flow cytometer (BD Biosciences).

## Results

### Altered arginine sensing associated mTORC1 signaling in AD brains

Several reports showed increased mTORC1 activation in human AD and that modulation of this complex impacts AD-related phenotypes in AD animal models (An et al., 2003; Caccamo et al., 2010; Li et al., 2005; Spilman et al., 2010). Additionally, recent findings identified several amino acid sensors specific to L-arginine that directly activate mTORC1 signaling (Chantranupong et al., 2016; Rebsamen et al., 2015). We utilized a focused real-time quantitative reverse transcription PCR (qRT-PCR) array with 48 gene transcripts involved in mTOR complex signaling. We measured the mRNA samples extracted from human postmortem brain hippocampal tissues of AD patients and age-matched controls. Overall, we found 14 gene transcripts (13 increased vs. one decreased) differentially regulated in the AD compared to the control samples, meeting either a p-value less than 0.05 or absolute fold change (FC)-value above 1.5 (**Figure 1, A, B**). Volcano plot and multi-group plot showed *MTOR, DEPTOR, DEPDC5, RAPTOR, NPRL3, WDR59, and CASTOR1* as the most significantly increased gene transcripts (*p* < 0.05, FC ≥ 1.5) and *TFEB* as the most down-regulated gene transcript (*p* < 0.05, FC ≤ -1.5) among AD tissues (**Figure 1, C, D**). GATSL3, which encodes cellular arginine sensor for mTORC1 subunit 1 (*CASTOR1*), a cytosolic arginine sensor, showed the largest increase in AD samples (*p* = 0.004, FC = 5.5). In relation, we also found that the lysosomal arginine sensor, solute carrier family 38 member 9 (*SLC38A9*), was elevated in AD, albeit a p-value above 0.05 (*p* = 0.360, FC = 1.85). GATOR1 (GTPase-activating protein (GAP) activity toward Rags complex 1), a negative regulator of mTORC1, had three increased subunits (*DEPDC5, NPRL2, NPRL3*) in AD (*p* < 0.05 or FC ≥ 1.5). GATOR2 (GAP activity toward Rags complex 2), a positive regulator of mTORC1, showed increased subunits (*WDR59, MIOS, WDR24*) in AD (*p* < 0.05 or FC ≥ 1.5). *LAMTOR3*, a subunit of RAGULATOR that serves as a positive regulator of mTORC1, was significantly increased in AD (*p* = 0.020, FC = 1.5). Most notably, four out of five of the mTORC1 subunits (*DEPTOR, MTOR, RPTOR, MLST8*) were significantly increased in AD brains (*p* < 0.05). Finally, *TFEB* (transcription factor EB), a key transcription factor responsible for initiating lysosome biogenesis for autophagy, was significantly decreased in AD (*p* = 0.031, FC = -1.61). Overall, these data suggest AD brains showed uncoupled mTORC1 signaling associated with arginine sensing and impaired autophagy.

**Figure 1.**
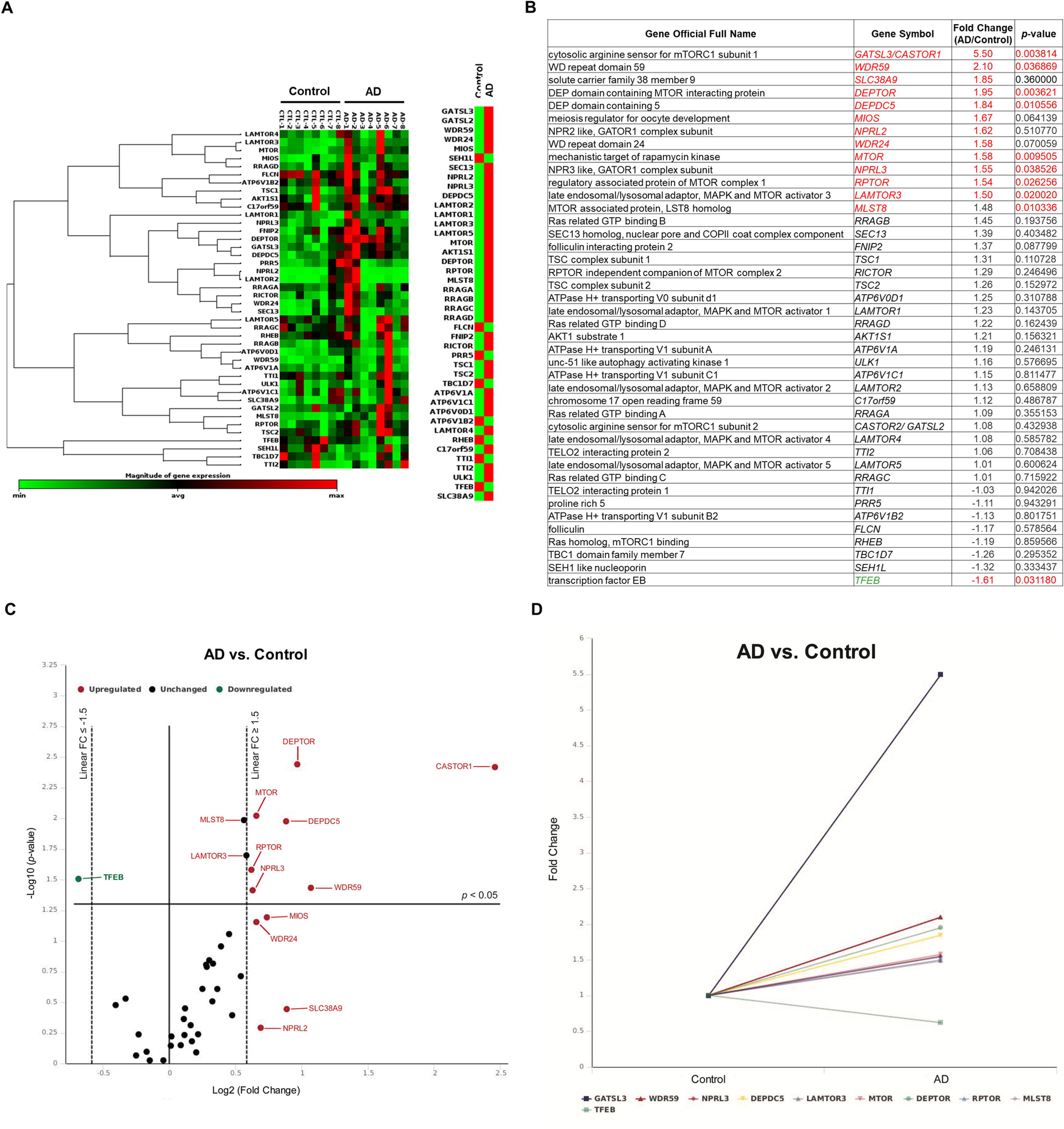
Alzheimer’s disease brains have increased expression of genes involved in arginine sensing-related mTORC1 signaling. A customized qRT-PCR array was performed using mRNA samples from postmortem hippocampus of Alzheimer’s disease (AD) patients and age-matched controls (Control). **A,** Gene expression heat map and cluster analysis revealed arginine sensing-related mTORC1 signaling gene transcripts among AD and control groups. **B,** Table of transcripts probed in the qRT-PCR array. Fold-change (FC) was calculated by comparing the AD and control groups. Red highlights indicate either p-value < 0.05 or absolute FC ≥ 1.5. Gene symbols highlighted in red and green indicate up-regulation and down-regulation, respectively. **C,** Volcano plot showed FC-value (Log2) versus p-value (-Log10) on the X and Y axes, respectively. Vertical dashes indicate FC-value = 1.5 or -1.5, and a horizontal solid line indicates p-value = 0.05. Up-regulated genes (p < 0.05 or FC ≥ 1.5) were annotated in red, down-regulated genes (p < 0.05 or FC ≤ -1.5) were in green, and unchanged genes (p > 0.05 or FC ≤ 1.5 and ≥ -1.5) were in black. **D,** Multi-group line plot specifically displayed the fold change of all statistically significant genes (p < 0.05) found in the AD group. Arginine Sensors: CASTOR1, SLC38A9; mTORC1 subunits: DEPTOR, MTOR, RPTOR, MLST8; GATOR1 subunits: DEPDC5, NPRL2, NPRL3; GATOR2 subunits: WDR59, MIOS, WDR24, SEC13, SEH1L; RAGULATOR subunits: LAMTOR1, LAMTOR2, LAMTOR3, LAMTOR4, LAMTOR5. Analyses and graphs were produced by Qiagen Online Data Analysis Center using Custom RT^2^ Profiler PCR Array. All genes were normalized to selected housekeeping genes. n = 8 per group. The p-values were calculated using an unpaired t-test of each gene’s replicate 2∧ (-Delta CT) values.

### Elevated expression of extracellular arginine receptor GPRC6A in AD brains

Given that several essential arginine sensing-related gene transcripts were increased in AD tissue, we measured the protein expression of GPRC6A, a putative extracellular arginine sensor/ receptor. We performed western blot analysis using whole cell lysate extracted from AD’s human postmortem brain cortex and age-matched controls. We measured tau deposition and found comparable levels of total monomeric tau between the AD and control tissues (*p* = 0.615, **Figure 2, A, B**, **Table 1**). However, total high molecular weight (HMW) tau was significantly enriched in the AD group (*p* = 0.003, **Figure 2, A, C**, **Table 1**). Furthermore, monomeric phospho-tau AT8 (pS202/Thr205) (*p* = 0.003, **Figure 2, A, D, Table 1**) along with its HMW (*p* = 0.001, **Figure 2, A, E, Table 1**) were both significantly increased in the AD group compared to the control group. Additionally, we measured phospho-tau at Ser214 because this epitope can become phosphorylated by mTORC1 substrate p70S6 Kinase 1 (S6K1) (Pei et al., 2006). Tau pS214 showed increased monomeric (*p* = 0.005, **Figure 2, A, F, Table 1**) and HMW (*p* = 0.002, **Figure 2, A, G, Table 1**) forms in AD, suggesting tau phosphorylation in AD were partly due to activated mTORC1 signaling. Most importantly, we found the AD brains showed elevated expression of GPRC6A (*p* = 0.024, **Figure 2, A, H, Table 1**) compared to the control group. These data potentially suggest tau accumulation in AD may promote the expression of GPRC6A for increased arginine sensing.

**Figure 2.**
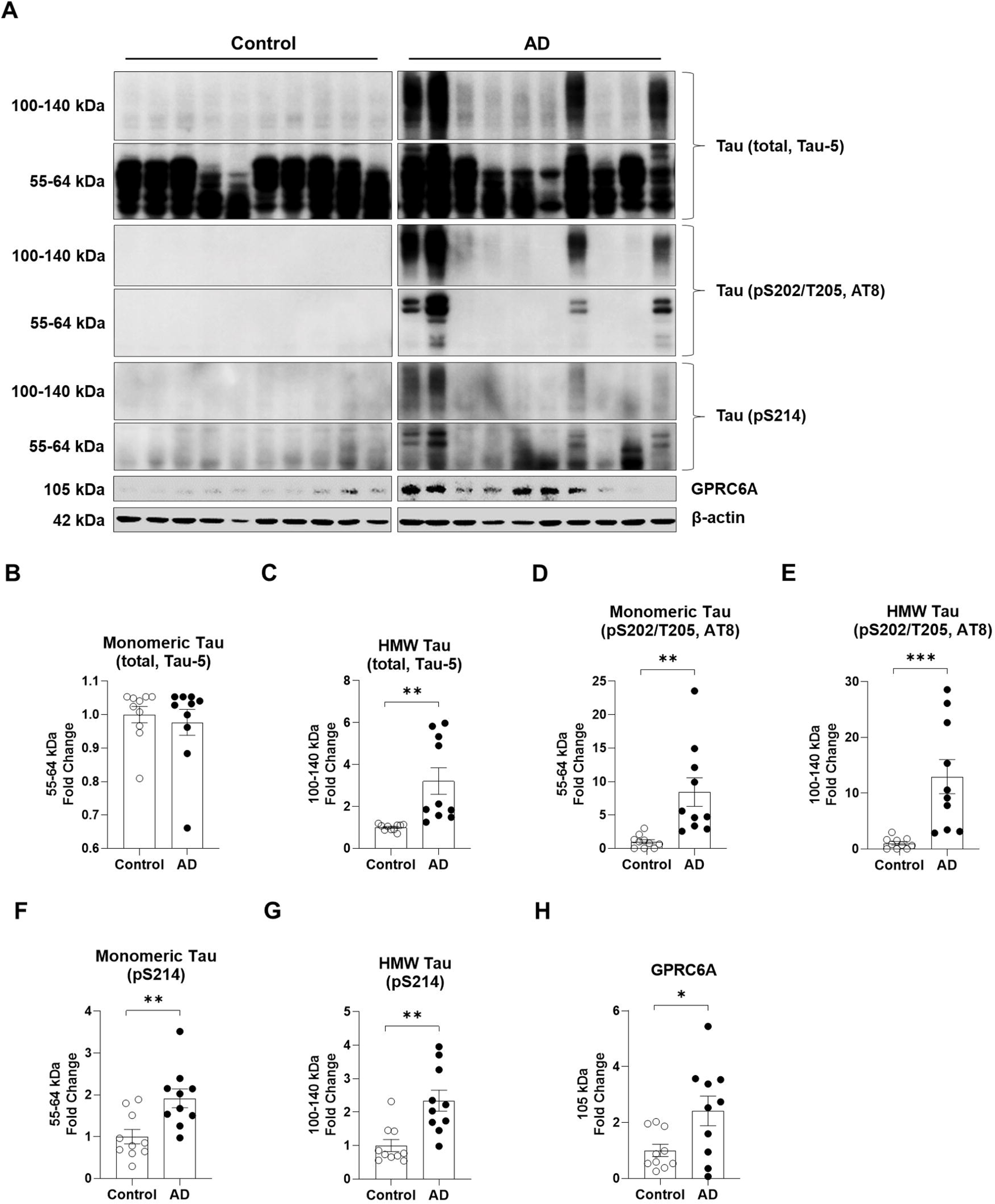
Alzheimer’s disease brains show increased tau and GPRC6A expression. Postmortem brain cortex samples from Alzheimer’s disease patients (AD) and age-matched controls (Control) were extracted with whole-cell lysate protein for western blot analysis. The AD group was normalized to the control group for comparison. **A,** Western blot images are presented for the Control and AD groups**. B-H,** Quantification of total monomeric tau (Tau-5, 55-64 kDa) (**B**), total high molecular weight (HMW) tau (Tau-5, 100-140 kDa) (**C**), monomeric phospho-tau (S202/T205, AT8, 55-64 kDa) (**D**), HMW phospho-tau (S202/T205, AT8, 100-140 kDa) (**E**), monomeric phospho-tau (S214, 55-64 kDa) (**F**), HMW phospho-tau (S214, 100-140 kDa) (**G**), and GPRC6A (**H**) are shown. n = 10 per group, * p < 0.05, ** p < 0.01, *** p < 0.001, unpaired t-test. Values represent mean ± SEM.

### Tau PS19 mouse brains show increased arginine anabolism and GPRC6A signaling

To understand the extent to which arginine metabolism and arginine sensing associated with GPRC6A signaling was disrupted in tauopathy mice, we extracted soluble protein from the posterior cortex of Tau PS19 mice and non-transgenic (Non-Tg) littermates for western blot analysis. The PS19 tau transgenic mice (*MAPT*, P301S) show neurofibrillary tangles at six months, remarkable neuronal loss at nine months, and cognitive deficit at ten months (Yoshiyama et al., 2007). In this current study, tau PS19 mice displayed increased levels of total tau (H150, *p* < 0.0001), phosphorylated tau at sites of Ser199/202 (*p* < 0.0001), Ser202/Thr205 (AT8, *p <* 0.0001), Ser396 (*p* < 0.0001), and Ser214 (*p* < 0.0001) compared to Non-Tg mice (**Figure 3, A, B, Table 1**). Argininosuccinate synthase 1 (ASS1) converts citrulline into argininosuccinate, which is further catalyzed into arginine by argininosuccinate lyase (ASL) (Haines et al., 2011). Although the expression of ASS1 (*p* = 0.420) did not change, the ASL (*p* < 0.0001) was increased in PS19 mice (**Figure 3, C-E, Table 1**), which may lead to an increased pool of arginine in the brain. We then measured the GPRC6A and found it was increased in Tau PS19 mice (*p* = 0.018, **Figure 3, C, F, Table 1**). Previous reports indicated that GPRC6A activation increased AKT and ERK1/2 signaling, and both act as positive upstream regulators for mTORC1 activation by inhibiting the TSC complex (Dangelmaier et al., 2014). We found the total level of AKT (*p* = 0.224) was unchanged, but the phospho-AKT (pT308, *p* = 0.048) was increased (**Figure 3, C, G, Table 1**). We also found that total ERK1/2 (*p* = 0.007) was elevated while the phospho-ERK1/2 (pT202/Y204, *p* = 0.931) did not change (**Figure 3, C, H, Table 1**). Collectively, these data indicate that tau overproduction in PS19 mice alters the expression of the anabolic arginine enzymes and increases GPRC6A expression and signaling.

**Figure 3.**
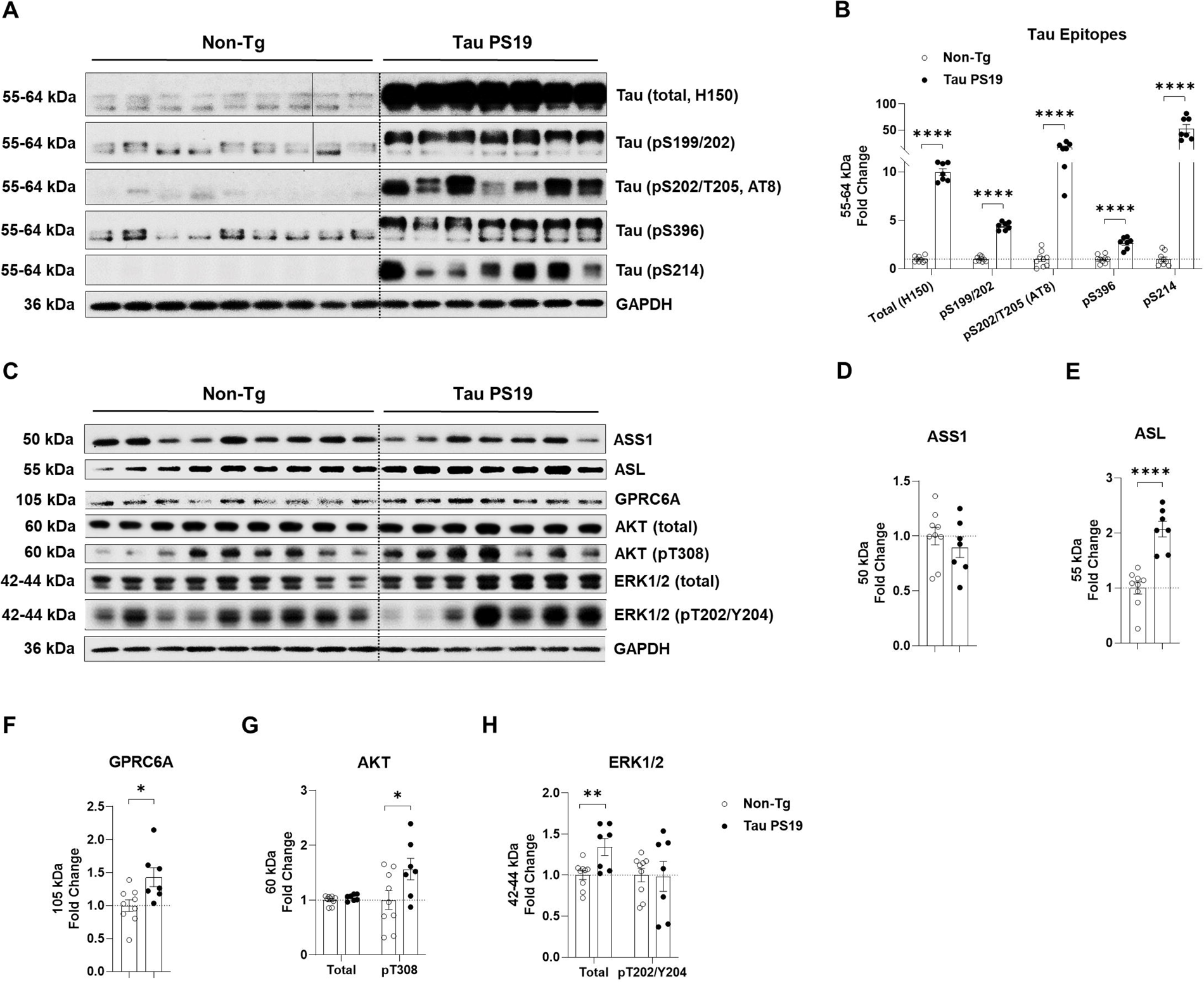
Tau PS19 mouse brains display elevated arginine anabolism and GPRC6A signaling. Mouse brain posterior cortex samples were homogenized and extracted with soluble protein for western blot. For comparison, the Tau PS19 mouse group was normalized to the non-transgenic littermates (Non-Tg) group. **A-B,** Tau PS19 mouse brains show increased total tau and phosphorylated tau epitopes. **A,** Western blot images are presented for total tau (H150) and several phospho-tau epitopes (pS199/202; Ser202/Thr205, AT8; Ser396; Ser214). **B,** Quantification analysis of A is shown. **C-H,** Tau PS19 mouse brains show elevated arginine-producing enzymes and GPRC6A signaling. **C,** Western blot images are shown for arginine-producing enzymes (ASS1, ASL), arginine receptor GPRC6A, AKT (total; pT308), and ERK1/2 (total; pT202/Y204). **D-H,** Quantification analysis of ASS1 (**D**), ASL (**E**), GPRC6A (**F**), AKT (**G**), and ERK1/2 (**H**) is shown. n = 7-9 mice per genotype, * p < 0.05, ** p < 0.01, **** p < 0.0001, unpaired t-test. Values are mean ± SEM.

### Tau PS19 mouse brains show increased mTORC1 signaling and decreased autophagy signaling

Next, we measured the total and phosphorylated proteins involved in mTORC1 activation in Tau PS19 mice. Although the total level of mTOR (*p* = 0.677) remained unchanged between the Tau PS19 mice and the Non-Tg littermates, phospho-mTOR (pS2448, *p* = 0.012) was increased in Tau PS19 mice (**Figure 4, A, B, Table 1**). Activated mTORC1 promotes protein synthesis through phosphorylation of crucial substrate effectors p70S6 Kinase 1 (S6K1), ribosomal protein S6 (S6), and eIF4E Binding Protein 1 (4E-BP1) (Liu and Sabatini, 2020). We found the total S6K1 (*p* = 0.713) did not change, while the phosphorylated S6K1 (pT421/S424, *p* = 0.041) was increased in Tau PS19 mice (**Figure 4, A, C, Table 1**). We also found that total S6 (*p* = 0.014) and phosphorylated S6 (pS235/236, *p* < 0.0001) were both increased in Tau PS19 mice (**Figure 4, A, D, Table 1**). Although the total level of 4E-BP1 (*p* = 0.882) remained unchanged, phosphorylation of 4E-BP1 (pT37/46, *p* = 0.014) was elevated in Tau PS19 mice (**Figure 4, A, E, Table 1**). These data suggest that tau PS19 mice show increased phosphorylation of key substrate effectors associated with mTORC1 activation.

**Figure 4.**
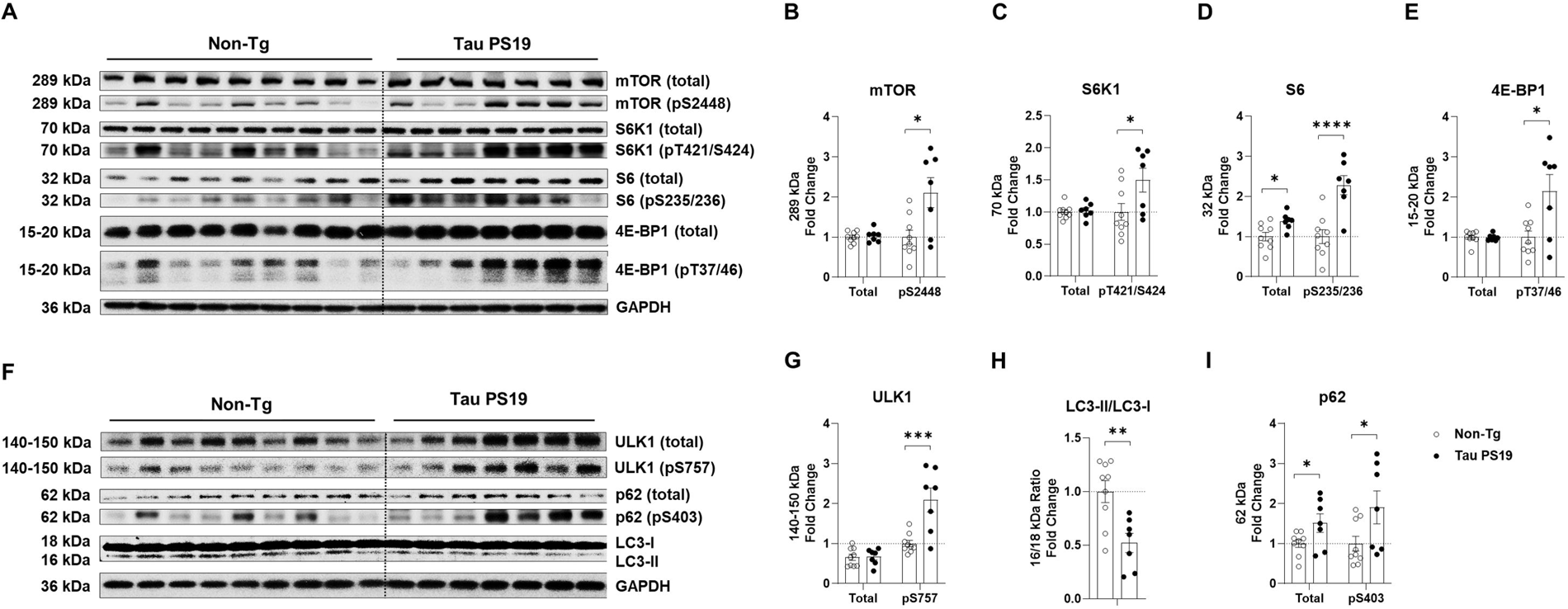
Tau PS19 mouse brains display activated mTORC1 signaling and reduced autophagy signaling. Mouse brain posterior cortex samples were homogenized and extracted for western blot analysis. Tau PS19 mouse group was normalized to the Non-Tg group for comparison. **A-C,** Tau PS19 mice show increased phosphorylation of the major mTORC1 subunit mTOR and several major mTORC1 downstream substrates (S6K1, S6, 4E-BP1). **A,** Western blot images of the total mTOR, S6K1, S6, 4E-BP1, and the associated phosphorylated mTOR (pS2448), S6K1 (pT421/S424), S6 (pS235/236), 4E-BP1 (pT37/46) are shown. **B-E,** Quantification analysis of A is shown for total and phosphorylated mTOR (**B**), S6K1 (**C**), S6 (**D**), and 4E-BP1 (**E**). **F-I,** Tau PS19 mice show increased phosphorylation of ULK1, p62, and decreased LC3-II/LC3-I ratio. **F,** Western blot images of total ULK1, p62, and the associated phosphorylated ULK1 (pS757), p62 (pS403), and LC3-I and LC3-II are shown. **G-I,** Quantification analysis of F is shown for total and phosphorylated ULK1 (**G**), p62 (**H**), and LC3-II/LC3-I ratio (**I**). n = 7-9 mice per genotype, * p < 0.05, ** p < 0.01, *** p < 0.001, **** p < 0.0001, unpaired t-test. Values are mean ± SEM.

In addition, other reports indicate that mTORC1 activation inhibits protein turnover by suppressing autophagy through the phosphorylation of Unc-51–like autophagy-activating kinase 1 (ULK1) (Kim et al., 2011). We found that total levels of ULK1 (*p* = 0.900) remained unchanged between Tau PS19 mice and non-Tg littermates, whereas phospho-ULK1 (pS757, *p* = 0.001) was increased (**Figure 4, F, G, Table 1**). We measured proteins associated with autophagic flux to determine if phosphorylated ULK1 inhibited the autophagy process. The formation and elongation of autophagosomes require the conversion of LC3-I into lipidated LC3-II to form the membrane. Therefore, LC3-II to LC3-I ratio typically indicates autophagic flux (Tanida et al., 2008). Compared to the Non-Tg littermates, Tau PS19 mice showed a decreased ratio of LC3-II to LC3-I (*p* = 0.004), suggesting slowed autophagic flux during tau neuropathology (**Figure 4, F, H, Table 1**). The final autophagolysosomes are formed when autophagosomes fuse with the lysosomes for degradation via the cargo adaptor protein sequestosome 1 (SQSTM1, p62) (Katsuragi et al., 2015). Thus, the accumulation of total p62 and its phosphorylation also indicate reduced autophagic flux (Ro et al., 2014). We found increased levels of total p62 (*p* = 0.043) and phospho-p62 (pS403, *p* = 0.047) in tau PS19 mice compared to the Non-Tg littermates (**Figure 4, F, I, Table 1**). Collectively, these data strongly suggest that tauopathy mouse brains show lysosomal dysfunction which may impair efficient autophagy flux.

### Tau PS19 mouse brains show increased arginine levels

We previously showed that mice with tauopathy (rTg4510 mice) displayed increased arginine levels in the CNS and that depletion of arginine by arginase1 overexpression impacted several components of mTORC1 and tau pathology (Hunt et al., 2015). Herein, we measured arginine levels in tau PS19 mice. We first measured total arginine abundance in the whole brain using whole-cell homogenates via liquid chromatography-mass spectrometry/mass spectrometry (LC-MS/MS) analysis. Arginine (*p* = 0.030) increased in Tau PS19 brains by 35% compared to Non-Tg littermates. However, ornithine (*p* = 0.296) and citrulline (*p* = 0.418) remained unchanged between the two groups (**Figure 5, A, Table 1**). We also assessed the extracellular level of arginine by collecting brain interstitial fluid using *in vivo* microdialysis. After allowing 18 hours of probe sampling, dialysates were measured by 10 hours for LC-MS/MS analysis (**Figure 5, B**). Over the 10-hour time window, Tau PS19 mice showed higher average baseline levels of interstitial arginine (*p* = 0.004) than Non-Tg littermates (**Figure 5, C, Table 1**). Western blot analysis also measured the total extracellular tau from the brain interstitial fluids. Interestingly, although we found monomeric tau probed by Tau-5 (*p* = 0.694) was unchanged between tau PS19 and Non-Tg mice, monomer tau measured by HT7 (*p* = 0.003) was increased in Tau PS19 mice (**Figure 5, D, E, Table 1**). Notably, HMW tau released into the brain interstitial fluid was increased in Tau PS19 mice, both probed by Tau-5 (*p* = 0.002) and HT-7 (*p* = 0.040) (**Figure 5, D, F, Table 1**). These data suggest that tauopathy mouse brains show increased arginine level in the presence of higher levels of interstitial tau.

**Figure 5.**
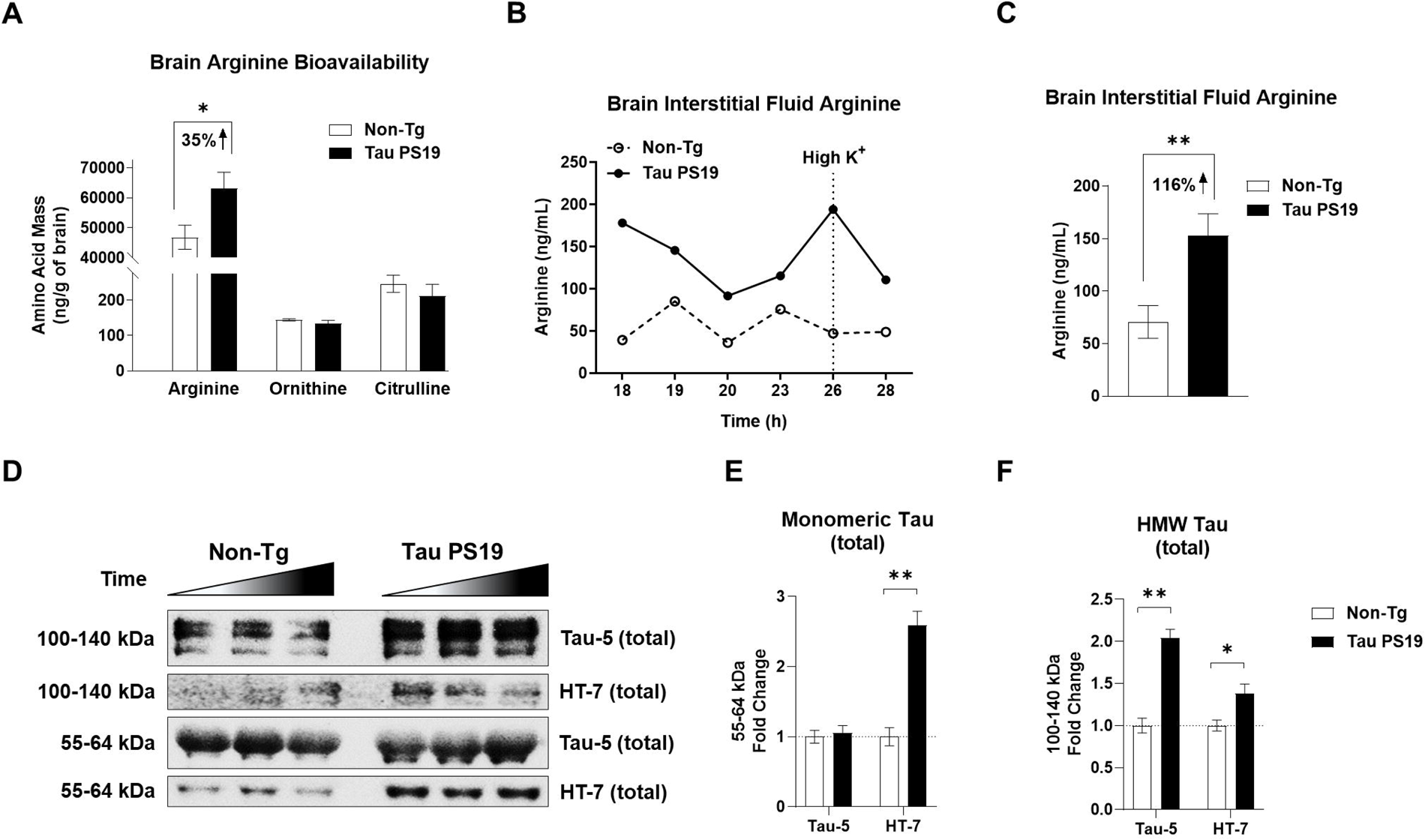
Tau PS19 mice harbor increased arginine level in the brain and brain interstitial fluid. Whole-brain homogenate and brain interstitial fluid from microdialysis were collected for LC-MS/MS measurement. **A,** Tau PS19 mouse brains show increased arginine and no change in ornithine and citrulline compared to the Non-Tg (n = 8 per genotype). **B-C,** Tau PS19 mice show higher levels of extracellular arginine in the brain’s interstitial fluid during the microdialysis (18-28 hours). **B,** Line plots of geometric mean for time points between 18 and 28 hours are shown. A high potassium pump was applied at a 26-hour time point. **C,** Quantification of arginine collected from brain interstitial of B is shown. **D-F,** Tau PS19 mice show increased total tau in brain interstitial fluids. Western blot images show total tau (Tau-5, HT-7) collected from brain interstitial fluid during microdialysis. **E-F,** Quantification analysis of monomeric tau (**E**) and high molecular weight (HMW) tau of Tau-5 and HT-7 is shown (**F**). The monomer tau of Tau-5 showed no change. Representative blot showing three-time points (n = 3 time points per genotype). * p < 0.05, ** p < 0.01, unpaired t-test. Values are mean ± SEM.

### Arginine supplementation activates mTORC1 signaling and promotes tau production in mouse primary cortical neurons

Given that arginine levels increase in the brain of the tau PS19 mice, along with elevated expression of GPRC6A and mTORC1 signaling, we sought to determine whether arginine directly promotes tau accumulation. We used arginine to activate mTORC1 in primary neurons to address this question. Arginine supplementation in primary mouse cortical neurons increased the phosphorylation of mTOR (pS2448, *p* = 0.027, **Figure 6, B**), S6 (pS235/236, *p* = 0.016, **Figure 6, C**), and 4E-BP1 (pT37/46, *p* = 0.033, **Figure 6, D**), while the total levels of mTOR (*p* = 0.359, **Figure 6, B**), S6 (*p* = 0.126, **Figure 6, C**), and 4E-BP1 (*p* = 0.554, **Figure 6, D**) remained unchanged (**Figure 6, A Table 1**). More importantly, we found that 1mM of arginine supplementation increased total tau level (Dako, *p* = 0.015) in primary neurons (**Figure 6, A, E Table 1**). Therefore, we found additional arginine stimulated mTORC1 signaling and promoted tau accumulation.

**Figure 6.**
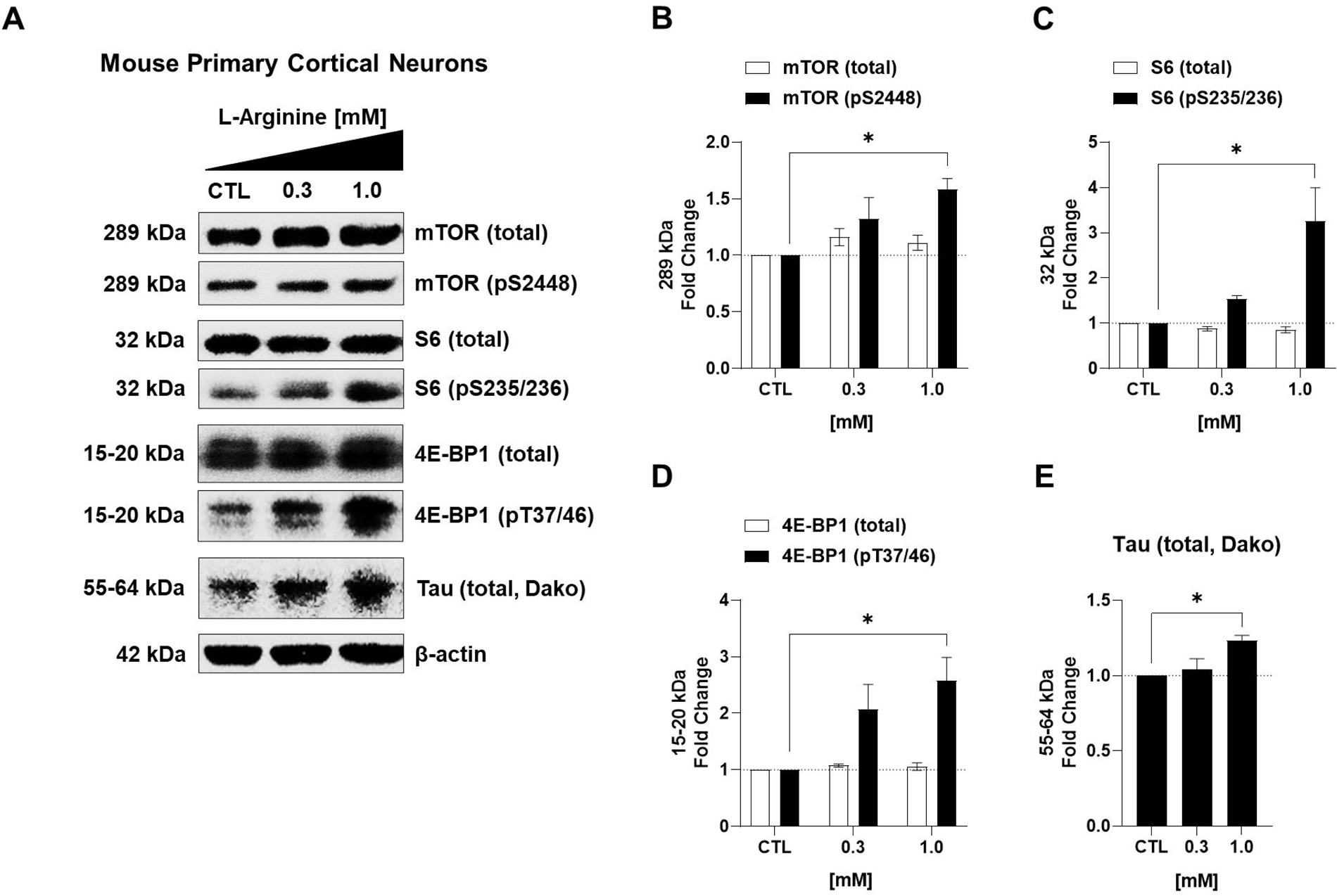
Arginine activates mTORC1 signaling and promotes tau accumulation in primary cortical neurons in mice. Arginine was supplemented in a concentration-dependent manner on the fourth-day culture of E18 wild-type mouse primary cortical neurons. The basal levels of arginine in the neurobasal medium (0.398 mM) served as the concentration for the control; two different concentrations of arginine (0.3 mM, 1 mM) were reconstituted into the medium. Data were normalized to the control for quantification analysis. After the treatment, cells were harvested and lysed as whole-cell lysate for western blot analysis. The arginine activated mTORC1 signaling, as measured by several downstream phosphorylated substrates. **A,** Representative western blot images of the total mTOR, S6, 4E-BP1, and the associated phosphorylated mTOR (pS2448), S6 (pS235/236), and 4E-BP1 (pT37/46), as well as total tau (Dako) are shown. **B-E,** Quantification analysis of western blots are shown for total and phosphorylated mTOR (**B**), S6 (**C**), 4E-BP1 (**D**), and total tau (**E**). n = 3 independent experiments, * p < 0.05, one-way ANOVA with Dunnett’s multiple-comparison test. Values are mean ± SEM.

### GPRC6A overexpression activates mTORC1 signaling and promotes tau accumulation and phosphorylation

Although we found that Tau PS19 mouse brains maintained a high level of arginine and elevated mTORC1 signaling *in vivo*, and arginine supplementation could activate mTORC1 and promote tau accumulation *in vitro*, we do not know if the arginine extracellular receptor GPRC6A plays a direct role in impacting mTORC1, autophagy, and tau accumulation. To determine if GPRC6A alters tau biology, we used HEK293T, which stably expresses a tetracycline-inducible tau (iHEK-Tau). We overexpressed the full-length membrane-bound *Gprc6a* fused with a Myc-tag or a green fluorescent protein (GFP) as a control, then analyzed it by western blot. We first measured the overexpressed protein products and found increased GPRC6A-Myc (*p* < 0.0001) and GFP (*p* = 0.007) expression (**Figure 7, A, B, Table 1**). GPRC6A is known to form dimers on the plasma membrane. We found monomeric and dimeric GPRC6A expression by western blot analysis (**Figure 7, A**) (Norskov-Lauritsen et al., 2015). We then measured the AKT-ERK signaling, comparing the GPRC6A-Myc and GFP control groups. We identified increased phosphorylation of AKT at T308 (*p* = 0.026) but not at S473 (*p* = 0.914) and no change in total AKT (*p* = 0.250) (**Figure 7, *A, C,* Table 1**) in GPRC6A-Myc treated cells. We also identified increased phosphorylation of ERK1/2 at T202/Y204 (*p* = 0.028) in the GPRC6A-Myc treated cells compared to GFP treated cells; however, the total level of ERK1/2 (*p* = 0.187) remained unchanged (**Figure 7, A, D, Table 1**). To assess the mTORC1 signaling, we found GPRC6A-Myc overexpression increased phosphorylation of mTOR (pS2448, *p* = 0.002) but not the total mTOR (*p* = 0.073) (**Figure 7, E, F, Table 1**). We also found increased phosphorylation of S6 at S235/236 (*p* = 0.005) and S240/244 (*p* = 0.005) in GPRC6A-Myc treated cells, while the total S6 (*p* = 0.629) remained unchanged compared to the GFP control (**Figure 7, E, G, Table 1**). In GPRC6A-Myc treated cells, 4E-BP1 phosphorylation at T37/46 (*p* = 0.028) and S65 (*p* = 0.031) increased, but no change was detected in the total levels (*p* = 0.065) (**Figure 7, E, H, Table 1**). Lastly, we sought to measure autophagy and tau production indicators during GPRC6A-Myc and GFP overexpression. We found GPRC6A-Myc overexpression increased phosphorylation of ULK1 (pS757, *p* = 0.036) but not in the total level (*p* = 0.215) compared to the GFP control (**Figure 7, I, J, Table 1**). The ratio of LC3-II/LC3-I was decreased in the GPRC6A-Myc group (*p* = 0.007, **Figure 7, I, K Table 1**), suggesting reduced autophagy flux. Importantly, we observed increased total tau (*p* = 0.040) and phosphorylated tau (pS214, *p* = 0.033) following GPRC6A-Myc overexpression (**Figure 7, I, L, Table 1**). In summary, this evidence strongly suggests that overexpressing GPRC6A elevates the AKT-ERK-mTORC1 signaling axis and impairs autophagy, thereby promoting the production and phosphorylation of tau in cellular models.

**Figure 7.**
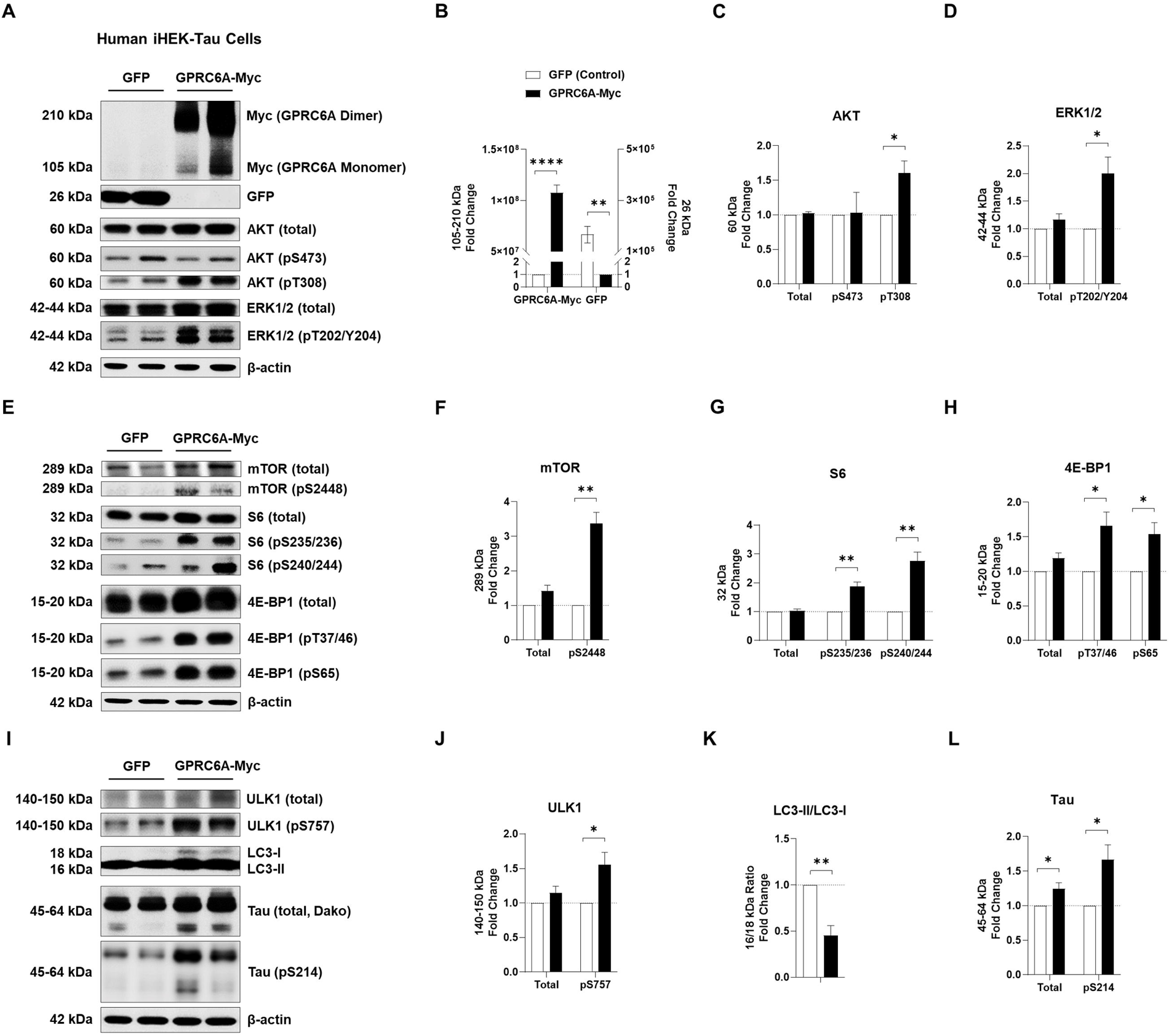
Overexpression of GPRC6A positively regulates mTORC1 signaling and increases tau in stably expressing cells. Tetracycline inducible human HEK293T cells overexpressing tau (iHEK-Tau) were transfected with plasmids to overexpress GPRC6A-Myc or GFP control. The GPRC6A-Myc group was normalized to the GFP control for quantification analysis. **A,** Representative western blot images of overexpressed proteins (GPRC6A-Myc, GFP) and AKT/ERK signaling-associated proteins are presented. **B-D,** Quantification analysis of western blots are shown for GPRC6A-Myc (**B**), GFP (**B**), total and phospho-AKT (pS473, pT308, **C**), and total and phospho-ERK (pT202/Y204, **D**). **E,** Representative western blot images of mTORC1 signaling-associated proteins are presented. **F-H,** Quantification analysis of E is shown for total and phospho-mTOR (pS2448, **F**), total and phospho-S6 (pS235/236, pS240/244, **G**), and total and phospho-4E-BP1 (pT37/46, pS65, **H**). **I,** Representative western blot images of autophagy-associated proteins and tau species are presented. **J-L,** Quantification analysis of I is shown for total and phospho-ULK1 (pS757, **J**), LC3-II/LC3-I ratio (**K**), and total and phospho-tau (pS214, **L**). n = 3 independent experiments, * p < 0.05, ** p < 0.01, **** p < 0.0001, unpaired t-test. Values are mean ± SEM.

### Genetic down-regulation of GPRC6A inhibits mTORC1 signaling and decreases tau expression

Given that genetic up-regulation of GPRC6A activated mTORC1 signaling and promoted tau production *in vitro*, we sought to determine if genetic repression of GPRC6A could potentially reverse this effect in human tau overexpressing cells. We transfected siRNAs against either human GPRC6A or a scramble sequence in human HeLa cells, stably overexpressing tau (C3HeLa Tau cells). Western blot analysis showed a decreased expression of GPRC6A (*p* = 0.002) upon transfection of siRNA against GPRC6A (**Figure 8, A, B, Table 1**). To assess the mTORC1 signaling, we found siRNA to GPRC6A decreased phosphorylation of mTOR (pS2448, *p* = 0.044) and the total mTOR (*p* = 0.002) (**Figure 8, A, C, Table 1**). We also found decreased phosphorylation of S6 at S235/236 (*p* < 0.0001) and total S6 (*p* = 0.001) in cells treated with siRNA to GPRC6A (**Figure 8, A, D, Table 1**). Additionally, phosphorylation of 4E-BP1 at T37/46 (*p* = 0.001) and total levels (*p* = 0.014) also decreased (**Figure 8, A, E, Table 1**) in response to siRNA to GPRC6a. To determine the downstream signaling of autophagy regulation, we measured ULK1. The GPRC6A siRNA transfection decreased the phosphorylation of ULK1 (pS757, *p* = 0.003) but not the total level (*p* = 0.446) (**Figure 8, A, F Table 1**). Furthermore, the ratio of LC3-II/LC3-I was increased in the GPRC6A siRNA-treated cells (*p* < 0.0001, **Figure 8, A, G, Table 1**). Remarkably, we observed a decrease in both total tau (*p* < 0.0001) and phosphorylated tau (Ps214, *p* < 0.0001) upon GPRC6A siRNA treatment (**Figure 8, A, H, Table 1**). Then, we performed a GPRC6A knockdown using the same siRNAs in the human iHEK-tau cells to validate these findings in another cell line. Indeed, GPRC6A expression decreased in iHEK tau cells compared to the scramble siRNA (*p* < 0.0001, **Figure 8, I, J, Table 1**). Similarly, the same critical substrates associated with mTORC1 and autophagy signaling were changed. siRNA to GPRC6A decreased total (*p* = 0.002) and phospho-mTOR (pS2448, *p* = 0.002) (**Figure 8, I, K, Table 1**), phosphor-S6 (pS235/236, *p* = 0.044) however total S6 (*p* = 0.818) remained unchanged (**Figure 8, I, L, Table 1**). Phospho-4E-BP1 (pT37/46, *p* = 0.001) decreased with GPRC6A repression; however, total levels were unchanged (*p* = 0.107) (**Figure 8, I, M, Table 1**). Additionally, GPRC6A repression decreased the phosphorylation of ULK1 (pS757, *p* = 0.039) but failed to modify its total levels (*p* = 0.374) (**Figure 8, I, N, Table 1**). The LC3-II/LC3-I ratio was increased during the GPRC6A knockdown (*p* = 0.025, **Figure 8, I, O, Table 1**). Finally, GPRC6A repression reduced the total tau (*p* = 0.007) and phospho-tau (pS214, *p* = 0.004) (**Figure 8, I, P, Table 1**). Collectively, these data strongly suggest that genetic down-regulation of GPRC6A inhibits mTORC1 signaling, activates autophagy, and reduces tau levels.

**Figure 8.**
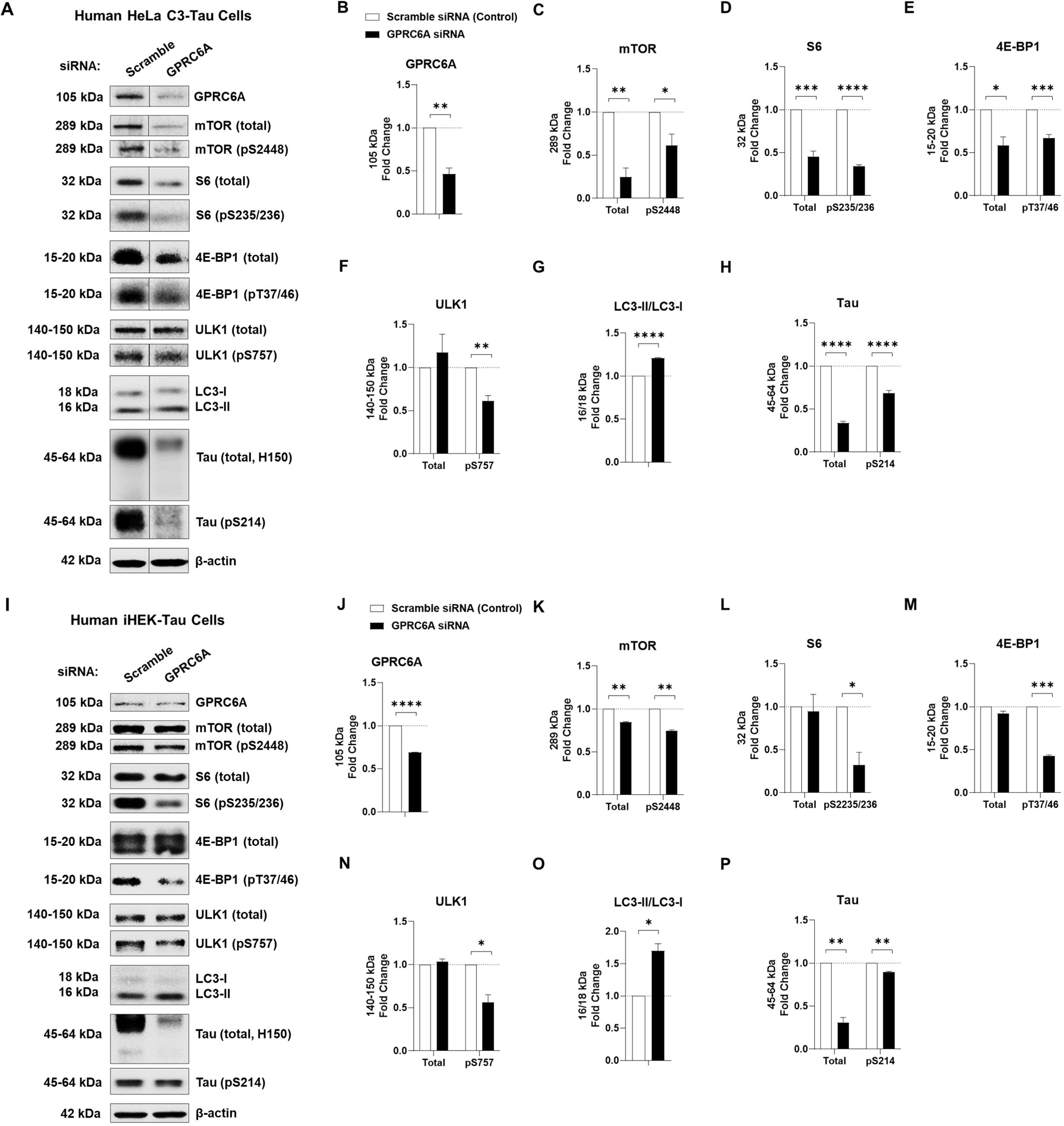
Down-regulation of GPRC6A negatively regulates mTORC1 signaling and decreases tau in stably expressing tau cells. **A-H**, Human HeLa cells stably overexpressing tau were transfected with siRNA to down-regulate the expression of GPRC6A. The scrambled siRNA served as the control. The siRNA group against GPRC6A was normalized to the siRNA control group for quantification analysis. **A,** Representative western blot images of GPRC6A, mTORC1-autophagy signaling associated proteins, and tau are presented. **B-H,** Quantification analysis of western blots are shown for GPRC6A (**B**), total and phospho-mTOR (pS2448, **C**), total and phospho-S6 (pS235/236, **D**), total and phospho-4E-BP1 (pT37/46, **E**), total and phospho-ULK1 (pS757, **F**), LC3-II/LC3-I ratio (**G**), and total and phospho-tau (pS214, **H**). n = 3 independent experiments. **I-P,** Tetracycline inducible human HEK293T cells overexpressing tau (iHEK-Tau) were transfected with either siRNA against GPRC6A or a scrambled target. The scrambled siRNA served as the control. The GPRC6A siRNA was normalized to the scramble siRNA control for quantification analysis. **I,** Representative western blot images of GPRC6A, mTORC1-autophagy signaling associated proteins, and tau are presented. **J-P,** Quantification analysis of I is shown for GPRC6A (**J**), total and phospho-mTOR (pS2448, **K**), total and phospho-S6 (pS235/236, **L**), total and phospho-4E-BP1 (pT37/46, **M**), total and phospho-ULK1 (pS757, **N**), LC3-II/LC3-I ratio (**O**), and total and phospho-tau (pS214, **P**). n = 2 independent experiments, * p < 0.05, ** p < 0.01, *** p < 0.001, **** p < 0.0001, unpaired t-test. Values are mean ± SEM.

### The recombinant adeno-assocaited virus delivery of shRNA against mouse Gprc6a decreases tau in PS19 mouse brains

We sought to determine if genetic repression of GPRC6A in the brains of mice impacts tau load. First, we tested two shRNA constructs against mouse *Gprc6a* by transfecting in mouse N2a cells. Western blot analysis showed that shGprc6a-2 (*p* = 0.007) decreased GPRC6A expression compared to the shScramble-GFP control-treated cells, while the shGprc6a-1 was not (*p* = 0.110) (**Figure 9, A, B, Table 1**). We also found that shGprc6a-2 reduced the total mTOR (*p* = 0.045) and the phosphorylated mTOR (pS2448, *p* = 0.039) (**Figure 9, A, C, Table 1**); however, the shGprc6a-1 showed no change for total mTOR (*p* = 0.132) and phospho-mTOR at S2448 (*p* = 0.109) (**Figure 9, A, C, Table 1**). Furthermore, cells treated with the shGprc6a-2 decreased the phosphorylation of S6 at S235/236 (*p* = 0.005) and S240/244 (*p* = 0.031), while the total S6 remained unchanged (*p* = 0.398) (**Figure 9, A, D, Table 1**). Cells treated with shGprc6a-1 failed to decrease the total S6 (*p* = 0.998), phospho-S6 of S235/236 (*p* = 0.092), and phospho-S6 S240/244 (*p* = 0.475) (**Figure 9, A, D, Table 1**). Therefore, the shGprc6a-2 shRNA construct was validated and packaged into the rAAV vector for mouse studies. The shScramble construct was also packaged into the same rAAV vector as the control.

**Figure 9.**
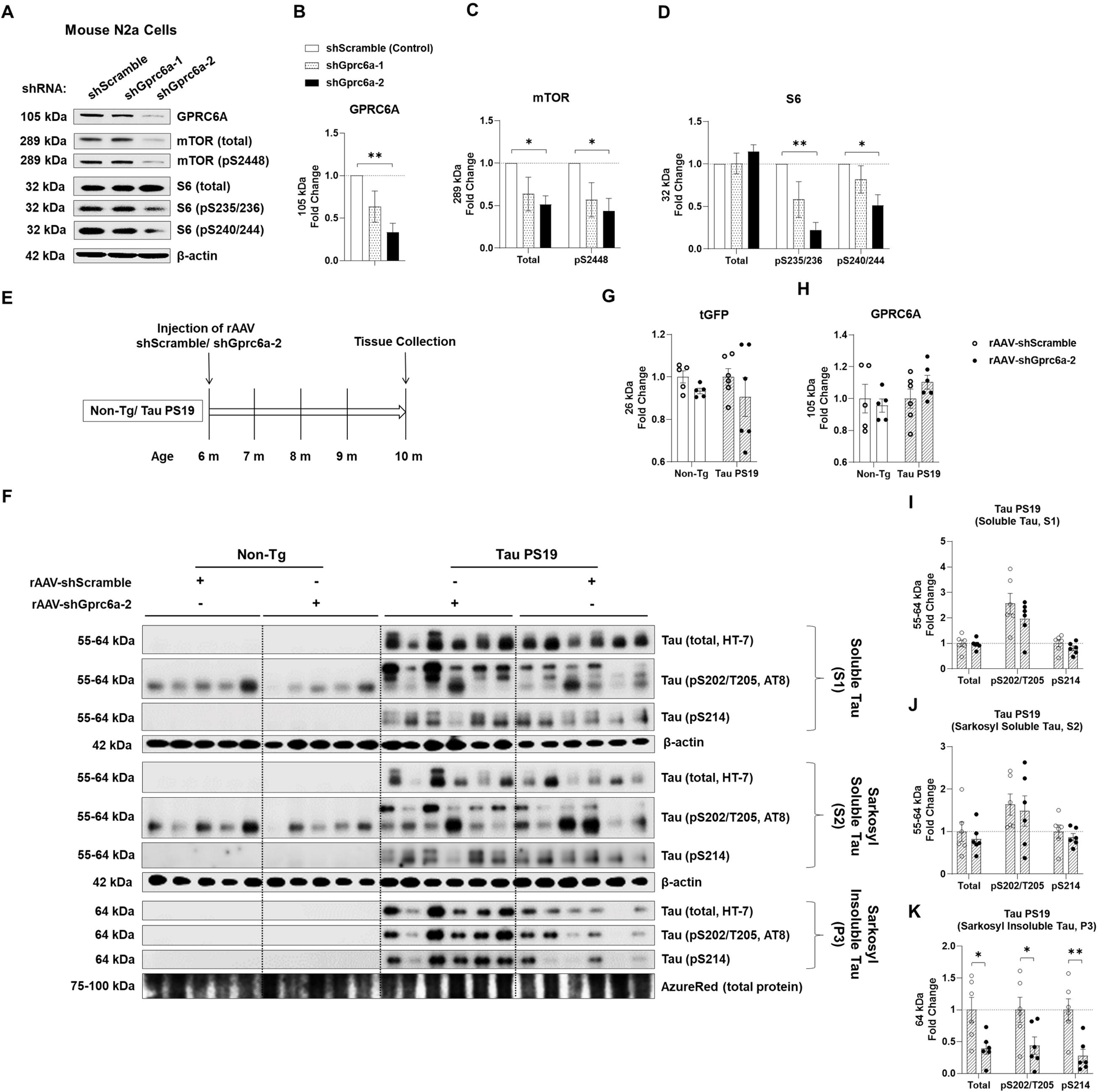
rAAV delivered shRNA against Gprc6a decreases sarkosyl insoluble tau in Tau PS19 mouse brains. **A-D**, Mouse N2a cells were transfected with two different shRNA against endogenous mouse Gprc6a and a scrambled sequence as the control. The two shRNA groups against Gprc6a (shGprc6a-1, shGprc6a-2) were normalized to the shRNA control group (shScramble) for quantification analysis. **A,** Representative western blot images of GPRC6A, total and phosphorylated mTOR, and S6 are shown. **B-D,** Quantification analysis of western blots for (**A**) is shown for GPRC6A (**B**), total and phospho-mTOR (pS2448, **C**), and total and phospho-S6 (pS235/236, pS240/244, **D**). n = 4 independent experiments. one-way ANOVA with Dunnett’s multiple-comparison test. **E,** Shows a schematic timeline diagram of the two rAAV constructs encoded with either shScramble or shGprc6a-2 injected into the brains of 6-month-old Non-Tg and Tau PS19. Four months post-incubation, brain tissue was harvested from 10-month-old mice and subjected to sequential sarkosyl-based protein extraction for western blot analysis. **F,** Western blot images of turbo green fluorescent protein (tGFP), GPRC6A, and total and phospho-tau are represented. **G-H,** Quantification analysis of F is shown for tGFP (**G**), GPRC6A (**H**) from the S1 soluble fraction and P1 membrane fraction of Non-Tg and Tau PS19 mice (n = 5-6 mice per group from Non-Tg and Tau PS19 mice). Two-way ANOVA followed by pairwise comparisons. **I-K,** Quantification analysis of F is shown for total tau (HT-7), phospho-tau at S202/T205 (AT8), and phospho-tau at S214 from protein fractions of soluble tau (S1, **I**), sarkosyl soluble tau (S2, **J**), and sarkosyl insoluble tau (P3, **K**) in Tau PS19 mice. n = 6 mice per group from Tau PS19 mice. Unpaired t-test. * p < 0.05, ** p < 0.01, Values are mean ± SEM.

Next, we injected the rAAV-shScramble-GFP and rAAV-shGprc6a-2 into the brains of Tau PS19 and their Non-Tg littermates at six months of age. After four months of viral incubation, we collected mouse brain tissues and performed protein extraction for tau using detergent-based sarkosyl fractionation measured by western blot analysis (**Figure 9, E**). To assess the rAAV transduction efficiency, we first measured the shared reporter protein turbo GFP (tGFP) expressed by both rAAV-shScramble-GFP and rAAV-shGprc6a-2. We found similar levels of tGFP expression in the total soluble fraction among all groups and had no overall change (*p* = 0.537) (**Figure 9, F, G, Table 1**). We then measured GPRC6A expression in the total membrane fraction but found that rAAV-shScramble and rAAV-shGprc6a-2 injection did not change the total brain GPRC6A level in Non-Tg and tau PS19 mice (*p* = 0.627) (**Figure 9, F, H, Table 1**). It is likely that the limited distribution of rAAV shRNA-GPRC6a expression in neurons was insufficient to detect a reduction, mainly when non-targeting cell types may also express the receptor. Nonetheless, we measured the tau levels in different fractions of protein extracts generated during the sarkosyl soluble and insoluble preparation. In the S1 fraction (non-detergent) soluble tau (total), we found that rAAV-shGprc6a-2 did not change the total tau levels (HT7, *p* = 0.814), phospho-tau at S202/T205 (AT8, *p* = 0.256), and phospho-tau at S214 (*p* = 0.261) compared to the rAAV-shScramble injected group (**Figure 9, F, I, Table 1**). In the S2 fraction (detergent sarkosyl soluble), we did not detect any changes in the total tau (HT-7, *p* = 0.533), phospho-tau S202/T205 (AT8, *p* = 0.733), and phospho-tau S214 (*p* = 0.485) either (**Figure 9, F, J, Table 1**). Interestingly, we did find that rAAV-shGprc6a-2 injection mice showed a reduction in total tau (HT-7, *p* = 0.018), phospho-tau S202/T205 (AT8, *p* = 0.041), and phospho-tau S214 (*p* = 0.005) in P3 fraction (sarkosyl insoluble tau) (**Figure 9, F, K, Table 1**). In summary, these data suggest that neuronal knockdown of mouse *Gprc6a* by rAAV delivery of shRNA decreases certain pools (sarkosyl insoluble) of tau in the PS19 mouse brains.

### Allosteric antagonism to GPRC6A mitigates tau oligomerization in vitro

Next, we used an allosteric antagonist to determine if GPRC6A inhibition could pharmacologically reduce tau expression. Cpd47661 was identified and characterized as a lead allosteric antagonist against GPRC6A (Gloriam et al., 2011). We previously established a novel mouse neuronal N2a split superfolder GFP-Tau (N2a-ssGT) cell line to stably express two human tau monomers individually fused with a split GFP portion. Tau oligomerization can be visualized and quantified upon complementing the two split GFP portions. We first validated the split GFP-Tau constructs by transfection in N2a naïve cells. The individual transfection of GFP10C (1-212 a.a.)-Tau or GFP11C (213-228 a.a.)-tau failed to produce any GFP signal; however, only the co-transfection of both constructs produced green fluorescence due to a complete GFP complementation (**Figure 10, A**). Following the transient transfection experiment, we created a stable monoclonal cell line using antibiotic selection (**Figure 10, A**). Compared to the N2a naïve cells, N2a-ssGT maintained a stable high percentage of GFP-positive cells at 55.3% (**Figure 10, B**). Therefore, a split GFP-Tau oligomerization assay was performed using this novel cell line to measure the effect of the GPRC6A allosteric antagonist in tau oligomerization. N2a-ssGT cells were incubated with Cpd47661 at 3 µM, 10 µM, and 30 µM, while the DMSO was used as the control (**Figure 10, C**). Post-treatment, cells were imaged for GFP fluorescence and harvested for flow cytometry analysis. We found Cpd47661 decreased GFP positive percentage at 3 µM (*p* = 0.012), 10 µM (*p* < 0.0001), and 30 µM (*p* < 0.0001) (**Figure 10, D, Table 1**). We also found Cpd47661 decreased green mean fluorescence intensity (MFI) at 3 µM (*p* < 0.0001), 10 µM (*p* < 0.0001), and 30 µM (*p* < 0.0001) (**Figure 10, E, Table 1**). A histogram analysis showed that each group’s average GFP positive percentage was decreased in a concentration-dependent manner (Control 55.4%, 3 µM 45.0%, 10 µM 38.4%, 30 µM 14.3%) (**Figure 10, F**). Overall, these data suggest that allosteric antagonism of GPRC6A reduced overall tau oligomerization *in vitro*.

**Figure 10.**
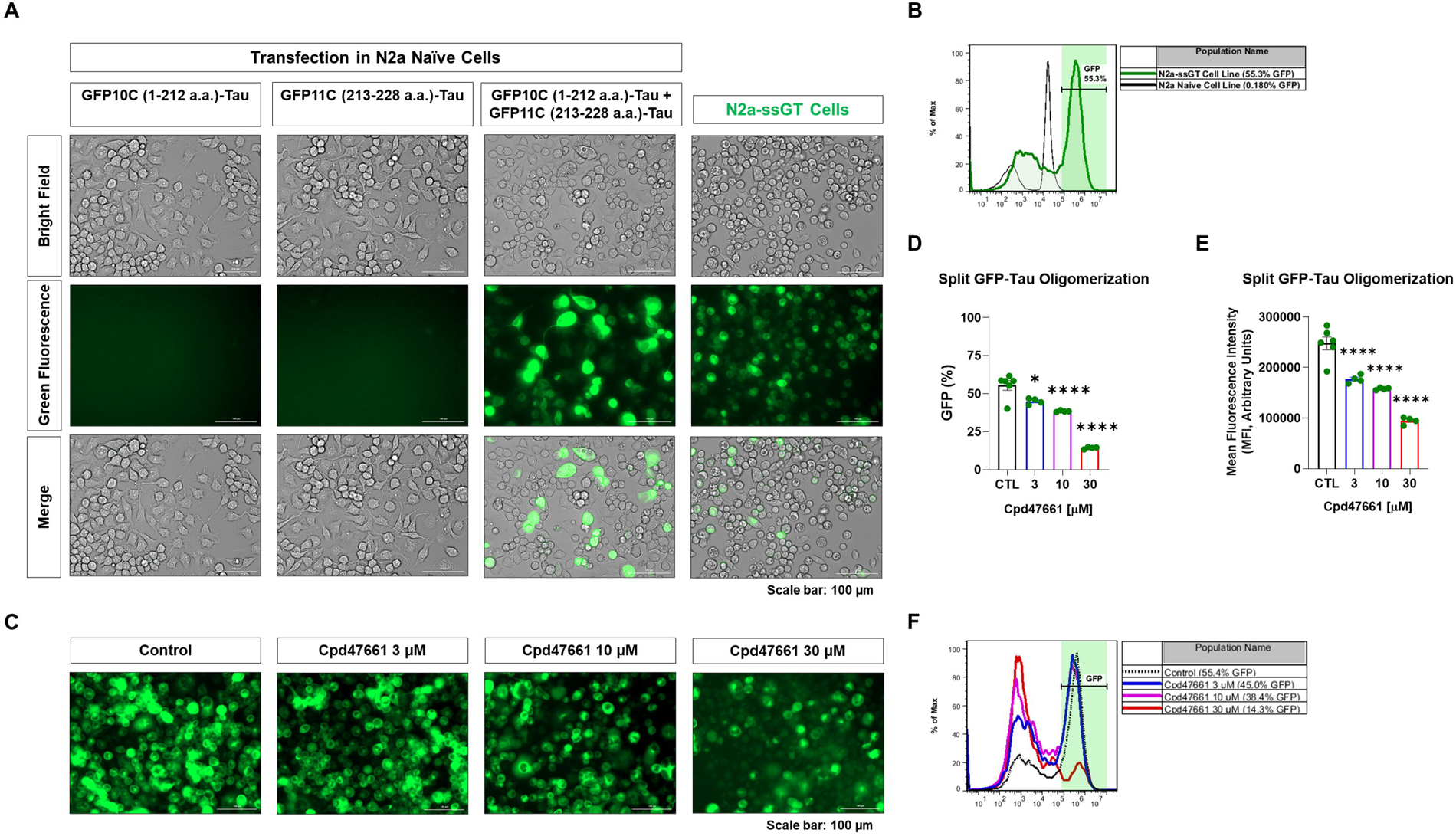
GPRC6A allosteric antagonist reduces tau accumulation and oligomerization in vitro. **A-B**, The N2a split superfolder GFP-Tau (N2a-ssGT) cell line was characterized as an *in vitro* model representing tau oligomerization by quantifying the GFP fluorescence. N2a-ssGT cells stably expressed two split GFP-Tau plasmids. **A,** Images of individual transfection of split GFP-Tau plasmids (GFP10C-Tau and GFP11C-Tau) and co-transfection in N2a naïve cells are represented. After cell selection with appropriate antibiotics, monoclonal cell line N2a-ssGT images are shown. **B,** The N2a-ssGT cell line was characterized by a separate GFP-positive cell population by flow cytometry. **C-F,** The GPRC6A allosteric antagonist Cpd47661 (3 µM, 10 µM, 30 µM) was added to N2a-ssGT cells. The DMSO as the vehicle was used as a control. All groups were subjected to imaging analysis for GFP fluorescence and flow cytometry analysis for GFP percentage. **C,** Representative images of N2a-ssGT cells incubated with different concentrations of Cpd47661 are shown. **D,** Flow cytometry analysis showed that the Cpd47661 decreased the GFP percentage in a concentration-dependent manner. **E,** Flow cytometry analysis showed that the Cpd47661 also decreased the GFP mean fluorescence intensity (MFI) in a concentration-dependent manner. **F,** An overlapped histogram showed that the GFP positive peak area decreased and the GFP negative peak area increased in a Cpd47661 concentration-dependent fashion. n = 4 independent experiments, * p < 0.05, **** p < 0.0001, one-way ANOVA with Dunnett’s multiple-comparison test. Values are mean ± SEM. The scale bar is 100 µm.

## Discussion

This study shows that tauopathies exhibit dysregulation of arginine-sensing proteins associated with mTORC1 signaling and reduced autophagy activity and that down-regulation of the arginine receptor GPRC6A can mitigate certain tau species. These findings are based on the following experimental evidence: First, we found that AD brains displayed elevated gene transcripts for mTORC1 signaling and intracellular arginine sensors (CASTOR1 and SLC38A9) but also increased GPRC6A expression at the protein level. Next, we discovered that mouse models of tauopathy showed increased total arginine level in the brains, coupled with anabolic enzymes for arginine production and signaling through GPRC6A and mTORC1. Finally, genetic manipulation of GPRC6A *in vitro* and *in vivo* modulated mTORC1 and autophagy signaling, as well as tau phenotypes. All of these strongly suggest that GPRC6A could be used as a therapeutic target for tauopathies or proteinopathies.

Previously, we showed that neuronal arginine depletion via ARG1 overexpression in rTg4510 tau transgenic mice reduced mTORC1 signaling and tau neuropathology (Hunt et al., 2015). We also found that ARG1 deficiency in brain myeloid cells of APP Tg2576 mice impaired Ragulator-Rag complex components involved in arginine sensing through mTORC1 signaling (Ma et al., 2021b). These findings indicate a link between arginine metabolism, abundance (repletion versus depletion), and signaling, which converges onto the mTORC1 pathway. We uncovered several signaling and functional outcomes by increasing or repressing GPRC6A. Therefore, we proposed a hypothetical model illustrating how tau aggregates could impair arginine metabolism or increase production, which may lead to increased arginine signaling associated with mTORC1 activation (**Figure 11, A**). Aberrant arginine sensing creates a feed-forward cycle of disease progression that may also be exaggerated in other chronic diseases, including cancer, diabetes, and the aging process (Liu and Sabatini, 2020). Despite some of the controversy surrounding different binding ligands, peripheral GPRC6A has been associated with osteocalcin-mediated endocrine networks, including testosterone signaling (Pi et al., 2015), bone mineralization (Pi et al., 2008; Pi et al., 2016), insulin secretion (Pi and Quarles, 2012b), as well as calcium-induced inflammation (Quandt et al., 2015; Rossol et al., 2012) and prostate cancer (Pi and Quarles, 2012a; Ye et al., 2017; Ye et al., 2019). Although GPRC6A is associated with the endocrine network in the peripheral, little is known about its respective role in the central nervous system, particularly during proteinopathies, neurodegeneration, and aging, likely due to its low expression in the CNS (Clemmensen et al., 2014; Jorgensen and Brauner-Osborne, 2020; Pi et al., 2017). However, it is still conceivable that cell specific low abundance proteins have a significant functional or pathological impact during neuronal challenges. Our current study shows that genetic knockdown of GPRC6a in PS19 with rAAV-shRNA reduces sarkosyl insoluble tau.

**Figure 11.**
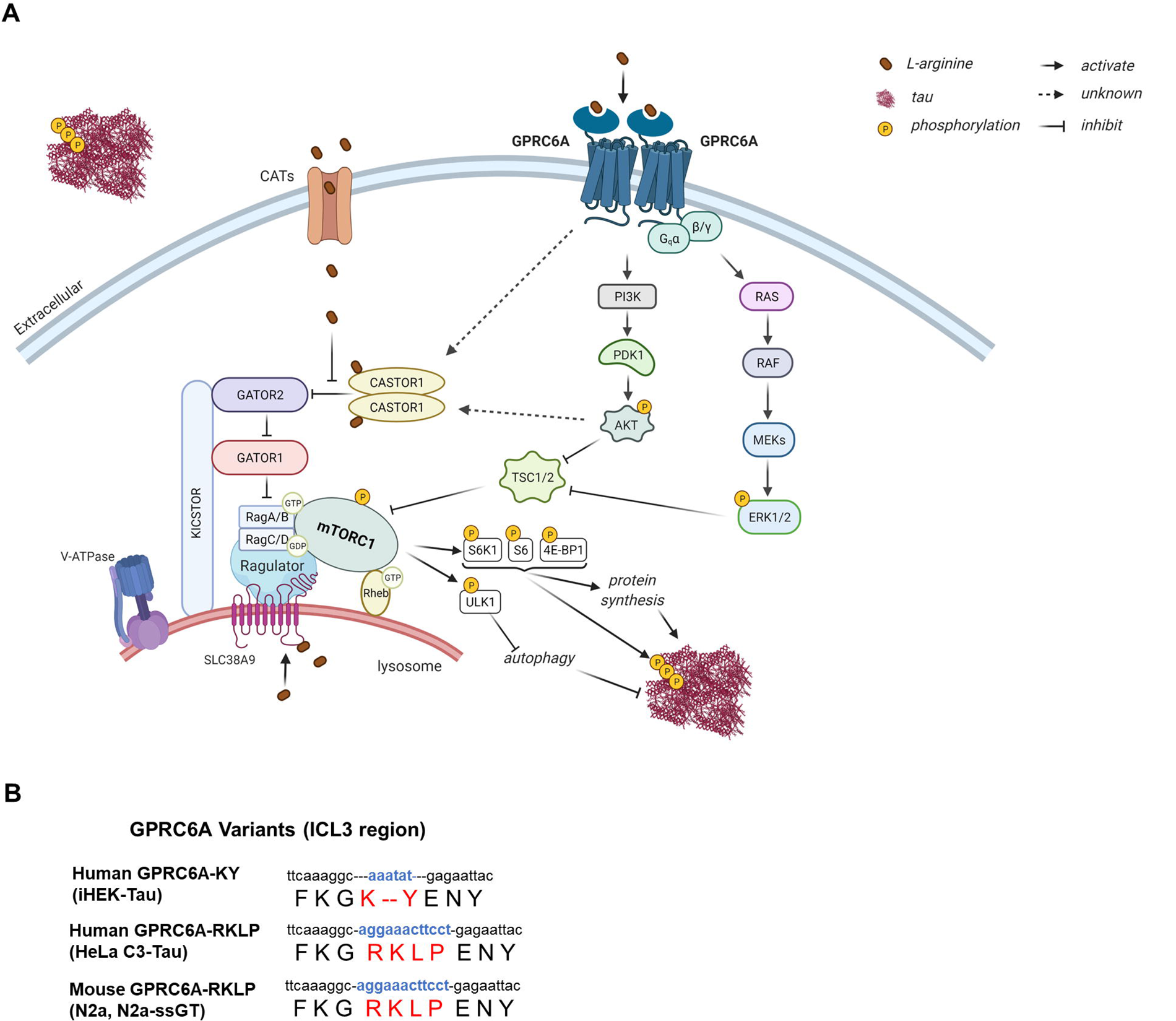
Hypothetical schematic diagram of arginine sensing associated mTORC1 signaling during tauopathies. **A**, Tau pathology impairs arginine metabolism, leading to arginine accumulation inside and outside neurons. Tau accumulation uncouples arginine sensing complexes to promote hyper mTORC1 activation. Positive and negative regulators of mTORC1 activation are listed as (±). Excess extracellular arginine can be taken into cells by cationic amino acid transporters (CATs) (+) to activate mTORC1 by binding to cytoplasmic arginine sensor CASTOR1 (-) and lysosomal arginine sensor SLC38A9 (+). Arginine can also bind to the extracellular arginine receptor GPRC6A (+) to induce mTORC1 activation, thus promoting protein synthesis and inhibiting autophagy. Complexes such as GATOR2 (+), GATOR1 (-), KICSTOR (-), RAGs (+), and Ragulator (+) coordinate the recruitment of mTORC1 to the lysosome membrane to interact with activated Rheb. Abbreviations include mTORC1: mechanistic target of rapamycin complex 1. CATs: cationic amino acid transporters. GPRC6A: G protein-coupled receptor (GPCR) family C, group 6, member A. PI3K: phosphoinositide 3-kinase. PDK1: 3-Phosphoinositide-dependent kinase 1. AKT: Protein kinase B (PKB). ERK1/2: extracellular-signal-regulated kinase 1/2. TSC1/2: tuberous sclerosis complex 1/2. CASTOR1: cellular arginine sensor for mTORC1 subunit 1. SLC38A9: solute carrier family 38 member 9. GATOR2: GTPase-activating protein (GAP) activity toward Rags complex 2. GATOR1: GAP activity toward Rags complex 1. KICSTOR: KPTN, ITFG2, C12orf66, and SZT2-containing regulator of mTORC1. V-ATPase: vacuolar-type H^+^-ATPase. Rag: Ras-related GTP binding protein. Ragulator: Lamtor1/2/3/4/5. Rheb: Ras homolog enriched in the brain. S6K1: p70 ribosomal S6 kinase. 4E-BP1: eukaryotic initiation factor 4E-binding protein 1. ULK1: Unc-51-like kinase 1. Created with BioRender.com **B,** Human and mouse cell lines used in the study were highlighted with the amino acid sequence of GPRC6A variants’ intracellular loop 3 (ICL3).

Increasing evidence suggests a metabolic role of GPRC6A expression; however, recent reports reveal that human GPRC6A harbors several major genetic variants comprised of a short insertion/ deletion (indel) sequence of *K..Y* in place of the long ancestral *RKLP* sequence located within the intracellular loop three (ICL3) region. The resulting SNP alters subcellular receptor distribution and Gq signaling (Jorgensen et al., 2017; Pi et al., 2017; Ye et al., 2017). The human GPRC6A-*K..Y* is retained intracellularly, while the full-length GPRC6A-*RKLP* is membrane-bound, as in other species like the mouse (**Figure 11, B**). The ICL3-*RKLP* allele is present in all other animal species and has a lower frequency in humans except for individuals of African descent (Ye et al., 2017). While the ICL3-*K..Y* dominates in humans and becomes the orthosteric reference allele (Pi et al., 2017), it is tempting to speculate that the ICL3-*RKLP* remains the ancestral allele that was evolutionarily conserved. Therefore, it is considered a polymorphism (rs386705086) primarily found in African ancestry populations. This ethnic difference indicates that the GPRC6A polymorphism may potentially influence disease risk associated with GPRC6A signaling. Importantly, we discovered that down-regulating GPRC6A in two crucial cell lines with different GPRC6A variants (iHEK-Tau harbors the GPRC6A-*K..Y* while HeLa C3-Tau harbors the GPRC6A-*RKLP* version) both decreased mTORC1 signaling and tau accumulation. These might suggest that specific signals of GPRC6A remain associated with mTORC1. Importantly, GPRC6A variants within the ICL3 region differed through signaling of D-*myo*-inositol monophosphate (IP_1_) accumulation assay and cell surface expression (Jorgensen et al., 2017). Conversely, we also found up-regulation of mouse GPRC6A-*RKLP* in iHEK-Tau cells, which endogenously harbors the *K..Y* allele, increased mTORC1 signaling and tau levels (**Figure 7**, **Figure 8**). However, it is unclear if the membrane-associated GPRC6A-*RKLP* variant promotes more mTORC1 or less signaling than the intracellular GPRC6A-*K..Y* variant, which could impact disease risk. Further work is warranted to understand the different signaling, interacting components, and functional aspects of different GPRC6A variants and their potential role in the CNS and peripheral systems. Currently, GPCR-targeted drugs remain the largest group in Food and Drug Administration (FDA)-approved drugs (34%) (Hauser et al., 2018). Progress in optimizing the lead compound of GPRC6A allosteric antagonist showed more selective candidates were developed (Johansson et al., 2015).

In conclusion, we found increased arginine pathways, molecular sensors, and mTORC1 signaling associated with tau biology in AD brains and tauopathy mouse models. This suggests a bi-directional link between arginine metabolism pathways and tau biology. Although the sequelae of arginine signaling dysfunction, tau biology, and hyper mTORC1 activation are unclear, their feed-forward cycling likely promotes disease progression. We posit that GPRC6A historically serves as an extracellular arginine sensor to tonically sense “nutrient (arginine) sufficient” states to maintain basal mTORC1 activation and balance the rate of autophagy. Genetic knockdown or pharmacological antagonism to GPRC6A signals “arginine deficiency,” thereby inhibiting mTORC1 signaling and increasing autophagy flux. Although further research is necessary, nutrient sensors, including GPRC6A, could be exploited as a novel therapeutic target for treating mTORopathies associated with nutrient-sensing dysfunction and proteostasis.

## REFERENCES

An, W. L., et al., 2003. Up-regulation of phosphorylated/activated p70 S6 kinase and its relationship to neurofibrillary pathology in Alzheimer’s disease. Am J Pathol. 163, 591–607.

Aurnhammer, C., et al., 2012. Universal real-time PCR for the detection and quantification of adeno-associated virus serotype 2-derived inverted terminal repeat sequences. Hum Gene Ther Methods. 23, 18–28.

Berger, Z., et al., 2007. Accumulation of pathological tau species and memory loss in a conditional model of tauopathy. J Neurosci. 27, 3650–62.

Caccamo, A., et al., 2010. Molecular interplay between mammalian target of rapamycin (mTOR), amyloid-beta, and Tau: effects on cognitive impairments. J Biol Chem. 285, 13107–20.

Chantranupong, L., et al., 2016. The CASTOR Proteins Are Arginine Sensors for the mTORC1 Pathway. Cell. 165, 153–164.

Clemmensen, C., et al., 2014. The GPCR, class C, group 6, subtype A (GPRC6A) receptor: from cloning to physiological function. Br J Pharmacol. 171, 1129–41.

Dangelmaier, C., et al., 2014. PDK1 selectively phosphorylates Thr(308) on Akt and contributes to human platelet functional responses. Thromb Haemost. 111, 508–17.

DeVos, S. L., et al., 2017. Tau reduction prevents neuronal loss and reverses pathological tau deposition and seeding in mice with tauopathy. Science Translational Medicine. 9.

Gloriam, D. E., et al., 2011. Chemogenomic discovery of allosteric antagonists at the GPRC6A receptor. Chem Biol. 18, 1489–98.

Graham, S. F., et al., 2015. Untargeted metabolomic analysis of human plasma indicates differentially affected polyamine and L-arginine metabolism in mild cognitive impairment subjects converting to Alzheimer’s disease. PLoS One. 10, e0119452.

Haines, R. J., et al., 2011. Argininosuccinate synthase: at the center of arginine metabolism. Int J Biochem Mol Biol. 2, 8–23.

Hauser, A. S., et al., 2018. Pharmacogenomics of GPCR Drug Targets. Cell. 172, 41–54 e19

Hunt, J. B., Jr., et al., 2015. Sustained Arginase 1 Expression Modulates Pathological Tau Deposits in a Mouse Model of Tauopathy. J Neurosci. 35, 14842–60.

Jacobsen, S. E., et al., 2013. Delineation of the GPRC6A receptor signaling pathways using a mammalian cell line stably expressing the receptor. J Pharmacol Exp Ther. 347, 298–309.

Johansson, H., et al., 2015. Selective Allosteric Antagonists for the G Protein-Coupled Receptor GPRC6A Based on the 2-Phenylindole Privileged Structure Scaffold. Journal of Medicinal Chemistry. 58, 8938–8951.

Jorgensen, C. V., Brauner-Osborne, H., 2020. Pharmacology and physiological function of the orphan GPRC6A receptor. Basic Clin Pharmacol Toxicol. 126 Suppl 6, 77–87.

Jorgensen, S., et al., 2017. Genetic Variations in the Human G Protein-coupled Receptor Class C, Group 6, Member A (GPRC6A) Control Cell Surface Expression and Function. J Biol Chem. 292, 1524–1534.

Kan, M. J., et al., 2015. Arginine deprivation and immune suppression in a mouse model of Alzheimer’s disease. J Neurosci. 35, 5969–82.

Katsuragi, Y., et al., 2015. p62/SQSTM1 functions as a signaling hub and an autophagy adaptor. FEBS J. 282, 4672–8.

Kim, J., et al., 2011. AMPK and mTOR regulate autophagy through direct phosphorylation of Ulk1. Nat Cell Biol. 13, 132–41.

La Joie, R., et al., 2020. Prospective longitudinal atrophy in Alzheimer’s disease correlates with the intensity and topography of baseline tau-PET. Sci Transl Med. 12.

Lasagna-Reeves, C. A., et al., 2016. Reduction of Nuak1 Decreases Tau and Reverses Phenotypes in a Tauopathy Mouse Model. Neuron. 92, 407–418.

Li, X., et al., 2005. Levels of mTOR and its downstream targets 4E-BP1, eEF2, and eEF2 kinase in relationships with tau in Alzheimer’s disease brain. FEBS J. 272, 4211–20.

Liu, G. Y., Sabatini, D. M., 2020. mTOR at the nexus of nutrition, growth, ageing and disease. Nat Rev Mol Cell Biol. 21, 183–203.

Liu, P., et al., 2014. Altered arginine metabolism in Alzheimer’s disease brains. Neurobiol Aging. 35, 1992–2003.

Ma, C., et al., 2021a. Myeloid Arginase 1 Insufficiency Exacerbates Amyloid-β Associated Neurodegenerative Pathways and Glial Signatures in a Mouse Model of Alzheimer’s Disease: A Targeted Transcriptome Analysis. Frontiers in Immunology. 12.

Ma, C., et al., 2021b. Arginase 1 Insufficiency Precipitates Amyloid-β Deposition and Hastens Behavioral Impairment in a Mouse Model of Amyloidosis. Frontiers in Immunology. 11.

Maeda, S., Mucke, L., 2016. Tau Phosphorylation-Much More than a Biomarker. Neuron. 92, 265–267.

Masters, C. L., et al., 2015. Alzheimer’s disease. Nat Rev Dis Primers. 1, 15056.

Morris, M., et al., 2011. The many faces of tau. Neuron. 70, 410–26.

Morrison, L. D., Kish, S. J., 1995. Brain polyamine levels are altered in Alzheimer’s disease. Neurosci Lett. 197, 5–8.

Mueed, Z., et al., 2018. Tau and mTOR: The Hotspots for Multifarious Diseases in Alzheimer’s Development. Front Neurosci. 12, 1017.

Norskov-Lauritsen, L., et al., 2015. N-glycosylation and disulfide bonding affects GPRC6A receptor expression, function, and dimerization. FEBS Lett. 589, 588–97.

Oddo, S., 2012. The role of mTOR signaling in Alzheimer disease. Front Biosci (Schol Ed). 4, 941–52.

Pei, J. J., et al., 2006. P70 S6 kinase mediates tau phosphorylation and synthesis. FEBS Lett. 580, 107–14.

Pei, J. J., Hugon, J., 2008. mTOR-dependent signalling in Alzheimer’s disease. J Cell Mol Med. 12, 2525–32.

Pi, M., et al., 2008. GPRC6A null mice exhibit osteopenia, feminization and metabolic syndrome. PLoS One. 3, e3858.

Pi, M., et al., 2015. Structural and Functional Evidence for Testosterone Activation of GPRC6A in Peripheral Tissues. Mol Endocrinol. 29, 1759–73.

Pi, M., et al., 2016. Evidence for Osteocalcin Binding and Activation of GPRC6A in beta-Cells. Endocrinology. 157, 1866–80.

Pi, M., et al., 2017. GPRC6A: Jack of all metabolism (or master of none). Mol Metab. 6, 185–193.

Pi, M., Quarles, L. D., 2012a. GPRC6A regulates prostate cancer progression. Prostate. 72, 399–409.

Pi, M., Quarles, L. D., 2012b. Multiligand specificity and wide tissue expression of GPRC6A reveals new endocrine networks. Endocrinology. 153, 2062–9.

Quandt, D., et al., 2015. GPRC6A mediates Alum-induced Nlrp3 inflammasome activation but limits Th2 type antibody responses. Sci Rep. 5, 16719.

Rebsamen, M., et al., 2015. SLC38A9 is a component of the lysosomal amino acid sensing machinery that controls mTORC1. Nature. 519, 477-+.

Ro, S. H., et al., 2014. Sestrin2 promotes Unc-51-like kinase 1 mediated phosphorylation of p62/sequestosome-1. FEBS J. 281, 3816–27.

Rossol, M., et al., 2012. Extracellular Ca2+ is a danger signal activating the NLRP3 inflammasome through G protein-coupled calcium sensing receptors. Nat Commun. 3, 1329.

Sandusky-Beltran, L. A., et al., 2019. Spermidine/spermine-N(1)-acetyltransferase ablation impacts tauopathy-induced polyamine stress response. Alzheimers Res Ther. 11, 58.

Sandusky-Beltran, L. A., et al., 2021. Aberrant AZIN2 and polyamine metabolism precipitates tau neuropathology. J Clin Invest. 131.

Spillantini, M. G., Goedert, M., 2013. Tau pathology and neurodegeneration. Lancet Neurol. 12, 609–22.

Spilman, P., et al., 2010. Inhibition of mTOR by rapamycin abolishes cognitive deficits and reduces amyloid-beta levels in a mouse model of Alzheimer’s disease. PLoS One. 5, e9979.

Tanida, I., et al., 2008. LC3 and Autophagy. Methods Mol Biol. 445, 77–88.

Vemula, P., et al., 2019. Altered brain arginine metabolism in a mouse model of tauopathy. Amino Acids. 51, 513–528.

Wang, C., et al., 2014. Targeting the mTOR signaling network for Alzheimer’s disease therapy. Mol Neurobiol. 49, 120–35.

Wellendorph, P., et al., 2005. Deorphanization of GPRC6A: a promiscuous L-alpha-amino acid receptor with preference for basic amino acids. Mol Pharmacol. 67, 589–97.

Wolfson, R. L., Sabatini, D. M., 2017. The Dawn of the Age of Amino Acid Sensors for the mTORC1 Pathway. Cell Metab. 26, 301–309.

Woo, J. A., et al., 2017. Loss of function CHCHD10 mutations in cytoplasmic TDP-43 accumulation and synaptic integrity. Nat Commun. 8, 15558.

Yamada, K., et al., 2014. Neuronal activity regulates extracellular tau in vivo. J Exp Med. 211, 387–93.

Ye, R., et al., 2017. CRISPR/Cas9 targeting of GPRC6A suppresses prostate cancer tumorigenesis in a human xenograft model. J Exp Clin Cancer Res. 36, 90.

Ye, R., et al., 2019. Human GPRC6A Mediates Testosterone-Induced Mitogen-Activated Protein Kinases and mTORC1 Signaling in Prostate Cancer Cells. Mol Pharmacol. 95, 563–572.

Yoshiyama, Y., et al., 2007. Synapse loss and microglial activation precede tangles in a P301S tauopathy mouse model. Neuron. 53, 337–51.

